# Synthetic lethality targeting LKB1 mutant and EGFR wild type human non-small cell lung cancer cells by glucose starvation and SGLT2 inhibition

**DOI:** 10.1101/622126

**Authors:** Yi Ren, Jiaqing Chen, Xiaofan Mo, Qiqi Yang, Peishi Chen, Guang Lu, Hayden Weng-Siong Tan, Juan Yi, Qiang Yu, You-Sun Kim, Karthik Mallilankaraman, Han-Ming Shen

## Abstract

In this study, we aimed to discover novel therapeutic approaches targeting non-small cell lung cancer (NSCLC) patients without EGFR mutation. First, we found that mutations of EGFR and LKB1 are mutually exclusive in NSCLC. EGFR-WT/LKB1-mutant cells are resistant to EGFR inhibitor erlotinib but are highly susceptible to glucose starvation or SGLT2 inhibitor canagliflozin. Mechanistically, in these cells, glucose starvation causes suppression of AMPK and induction of oxidative stress, leading to cell death. Finally, canagliflozin effectively reduces tumor growth of EGFR-WT/LKB1-mutant NSCLC cells in the mice xenograft model. Our data thus demonstrate that synthetic lethality can be achieved by glucose starvation or SGLT2 inhibition in EGFR-WT/LKB1-mutant NSCLC.

**SIGNIFICANCE:** At present, EGFR inhibitor-based targeted therapy can only benefit those non-small cell lung cancer (NSCLC) patients with EGFR mutation. Therefore, there is an urgent need to develop alternate targeted therapy for NSCLC with WT EGFR. In this study, we found that mutations of EGFR and LKB1 are mutually exclusive in NSCLC, and more importantly, synthetic lethality can be achieved in EGFR-WT/LKB1-mutant NSCLC cells with glucose starvation or SGLT2 inhibition. Since SGLT2 inhibitors such as canagliflozin are FDA-approved drugs for type II diabetes, our study thus points out a possibility of developing SGLT2 inhibitors as a targeted therapy for NSCLC patients with WT EGFR and mutant LKB1, which will benefit about 15-30% of NSCLC patients.

**HIGHLIGHTS:**

- EGFR and LKB1 mutations are mutually exclusive in NSCLC
- EGFR-WT/LKB1-mutant NSCLC cells are sensitive to cell death induced by glucose starvation and SGLT2 inhibition
- Glucose starvation suppresses AMPK activity in LKB1-mutant NSCLC cells
- SGLT2 inhibitor canagliflozin causes synthetic lethality in LKB1-mutant NSCLC cells

## INTRODUCTION

In the past two decades, one major breakthrough in lung cancer research is the discovery of the epidermal growth factor receptor (EGFR) mutations and successful development of specific EGFR tyrosine kinase inhibitors (TKIs) as a targeted therapy for non-small cell lung cancer (NSCLC) (Herbst et al., 2018). So far, three generations of EGFR TKIs have been developed and approved for the treatment of NSCLC with mutant EGFR. However, only a relatively small proportion of NSCLC patients were found with EGFR mutations (10% to 50%, depending on ethnicity) (Sharma et al., 2007). Therefore, the remaining large proportion of NSCLC patients with wild-type (WT) EGFR cannot benefit from these EGFR TKIs, indicating a huge gap in NSCLC treatment and an urgent need to develop novel targeted therapies that are applicable to NSCLC patients with WT EGFR.

The liver kinase B1 (LKB1) was originally identified as the tumor suppressor responsible for the inherited cancer disorder Peutz-Jeghers Syndrome (PJS) (Hemminki et al., 1998). The tumor suppressive functions of LKB1 has been extensively studied (Kullmann and Krahn, 2018; Momcilovic and Shackelford, 2015). LKB1 suppresses cell growth and cell proliferation when energy and nutrients are scarce, a process primarily achieved through the activation of the AMP-activated protein kinase (AMPK). The LKB1-AMPK axis inhibits the mechanistic target of rapamycin (mTOR) complex 1 (mTORC1) and activates autophagy via multiple pathways (Egan et al., 2011; Gwinn et al., 2008; Inoki et al., 2003; Kim et al., 2013; Kim et al., 2011). Moreover, AMPK phosphorylates and inactivates acetyl-CoA carboxylase (ACC) to inhibit fatty acid synthesis and promote fatty acid oxidation, therefore maintaining ATP and NADPH levels (Jeon et al., 2012; Li et al., 2018; Munday et al., 1988). Among many types of cancers involving LKB1, the implication of LKB1 in NSCLC appears to be particularly important. Loss-of-function mutations of LKB1 are also frequently detected in at least 15% to 30% of NSCLC patients (Ding et al., 2008; Gill et al., 2011; Ji et al., 2007). Functionally, it has been reported that in the presence of KRAS mutation, LKB1 mutation drastically increases tumor burden, and causes more frequent differentiation and metastasis (Ji et al., 2007). LKB1 mutation also promotes the development of lung adenocarcinoma to squamous cell carcinoma, leading to therapy resistance (Li et al., 2015). In contrast, several recent studies suggest that LKB1 inactivation is associated with the vulnerability of NSCLC cells to metabolic stresses, such as glucose starvation, matrix detachment, and the mitochondrial inhibitor phenformin (Jeon et al., 2012; Shackelford et al., 2013). Interestingly, LKB1 mutation was found to be closely associated with KRAS mutation in NSCLC (Ding et al., 2008; Ji et al., 2007; Makowski and Hayes, 2008), while KRAS mutation and EGFR mutation is known to be mutually exclusive in NSCLC (Kobayashi et al., 2005; Pao et al., 2005a). At present, the functional relationship between LKB1 and EGFR has not been studied.

Glucose is the most important source of energy for all organisms. Chemically, glucose is a polar molecule and it cannot permeate through the cell membrane by simple diffusion. The uptake of glucose is accomplished by glucose transporters. There are two major families of glucose transporters, *i.e.*, glucose transporters (GLUTs) and sodium-glucose co-transporters (SGLTs), distinguished by their mechanisms of action (Navale and Paranjape, 2016). Several GLUTs have been reported to be implicated in cancer. For example, GLUT1 is overexpressed in many types of cancers, probably induced by KRAS or BRAF mutations (Szablewski, 2013; Yun et al., 2009). Small molecule GLUT1 inhibitors have been described to selectively kill cancer cells *in vitro* (Chan et al., 2011; Liu et al., 2012; Shibuya et al., 2015). However, the ubiquitous expression of GLUT1 in normal cells may render these inhibitors inapplicable for clinical use. On the other hand, SGLTs are mainly expressed in intestine and kidney, especially for SGLT2, which is predominately expressed in the proximal tubule of the nephron and contributes to up to 90% of glucose reabsorption (Wright, 2001). Based on this, SGLT2 inhibitors, named gliflozins, have been successfully developed as the first line anti-diabetic drugs, including canagliflozin, dapagliflozin, and empagliflozin (Haas et al., 2014). Recently, SGLT2 has been shown as a diagnostic and therapeutic target for early-stage lung adenocarcinoma, and SGLT2 inhibitors greatly reduced lung tumor growth (Scafoglio et al., 2018). At this stage, it is not known whether the therapeutic effects of SGLT2 inhibitors are associated with the LKB1-AMPK pathway.

In this study, we first found that LKB1 mutation and EGFR mutation are mutually exclusive in human NSCLC patients. Moreover, those NSCLC cells with mutant LKB1 and WT EGFR are highly susceptible to cell death induced by glucose starvation, a process involving impaired AMPK activation and induction of oxidative stress. Finally, we found that SGLT2 inhibitors are capable of selectively killing NSCLC cells with mutant LKB1 and WT EGFR both *in vitro* and *in vivo*. Taken together, our results suggest that inhibition of glucose uptake, either by glucose deprivation or SGLT2 inhibition, in NSCLC cells that harbor LKB1 mutations results in synthetic lethality to these cells. These findings demonstrate the possibility of developing SGLT2 inhibitors as a novel targeted therapy for EGFR-WT NSCLC patients, thus filling a significant gap in lung cancer therapy.

## RESULTS

### EGFR and LKB1 Mutations Are Mutually Exclusive in NSCLC

The oncogenic EGFR and KRAS mutations are known to be mutually exclusive in NSCLC and in several other cancer types (Choughule et al., 2014; Fu et al., 2014; Kris et al., 2011; Shigematsu et al., 2005; Tam et al., 2006). Moreover, KRAS mutation is known to be negatively associated with the responsiveness of NSCLC to EGFR TKIs, serving as a negative biomarker for EGFR TKIs-based targeted therapies for NSCLC (Li et al., 2014b; Mao et al., 2010; Pao et al., 2005b). Interestingly, LKB1 mutations have been reported to be commonly co-occurred with KRAS mutations in NSCLC (Matsumoto et al., 2007; Schabath et al., 2016; Skoulidis et al., 2015). Thus, we hypothesized that a mutual exclusivity also exists between EGFR and LKB1 mutations. To prove this, we conducted a pooled analysis of EGFR, KRAS and LKB1 gene alterations using online database cBioPortal (http://www.cbioportal.org) (Cerami et al., 2012; Gao et al., 2013). There are 9 reported NSCLC studies with a total of 3,129 samples in 3,074 patients (Campbell et al., 2016; Cancer Genome Atlas Research, 2012, 2014; Ding et al., 2008; Imielinski et al., 2012; Jordan et al., 2017; Rizvi et al., 2018; Rizvi et al., 2015; Vavala et al., 2017). Consistent with previous studies, EGFR and KRAS mutations were mutually exclusive, whereas KRAS and LKB1 mutations were co-occurred in NSCLC (**Figures 1A and 1B, S1A and S1B**). One key finding from this analysis is that LKB1 mutation and EGFR mutation is mutually exclusive (**Figures 1A and 1B**). From a detailed analysis of these 9 reports (**Figure S1C**), we found that EGFR and LKB1 mutations are mutually exclusive in 8 of them, and the only negative result in Study #9 contains only 41 samples (Vavala et al., 2017). **Figure 1A** also revealed the types of mutations in EGFR and LKB1. For EGFR, the major types are amplification (in red), inframe mutation (in brown) and missense mutation (in dark green), while for LKB1, the major type is truncation (in black), being consistent with the expected functional implication in lung cancer. Based on the results from this set of analysis (**Figures S1C**), the percentage of LKB1 mutation in NSCLC ranged from 1.7% to 23%. The low percentage of LKB1 mutation in Study #8 was probably caused by the detection method used in the study (Cancer Genome Atlas Research, 2012).

**Figure 1.**
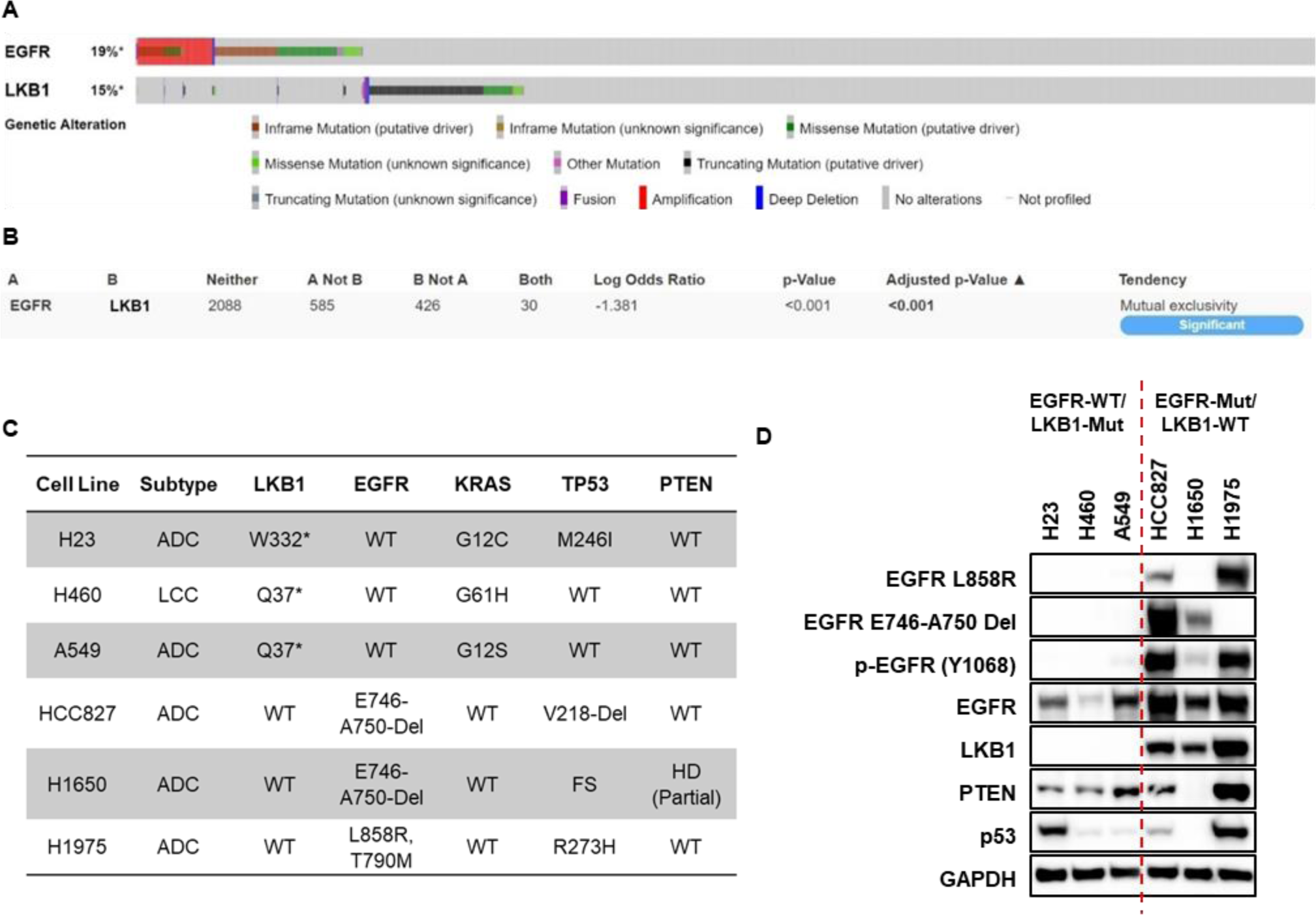
EGFR and LKB1 Mutations Are Mutually Exclusive in NSCLC. (A) The mutation spectrum of EGFR and LKB1 shows the discrete distribution of EGFR and LKB1 mutations in NSCLC. Nine NSCLC studies with 3,129 samples were pooled and analyzed using the cBioPortal database.
(B) The mutations of EGFR and LKB1 are mutually exclusive in NSCLC samples. Same sets of samples in (**A**) were tested for their mutual exclusivity using the cBioPortal database.
(C and D) The mutations of EGFR and LKB1/KRAS are mutually exclusive in NSCLC cell lines. **(C)** All information was obtained from COSMIC (https://cancer.sanger.ac.uk/cosmic) and ATCC (https://www.atcc.org/). ADC, adenocarcinoma; LCC, large cell carcinoma; *, non-sense mutation; WT, wild type; Del, deletion; FS, frame shift; HD, homozygous deletion; PM, promoter methylation. **(D)** Cells cultured under the normal condition were lysed. The protein expression was detected by immunoblotting using specific antibodies. See also Figure S1.

To further confirm the mutual exclusivity between EGFR and LKB1 mutations in NSCLC, we obtained 6 NSCLC cell lines from the American Type Culture Collection (ATCC) as listed in **Figure 1C**, and detected the expression of EGFR and LKB1 in these cells. Among them, 3 cell lines (H23, H460, and A549) are LKB1-mutant but EGFR-WT, whereas the other 3 cell lines (HCC827, H1650, and H1975) are LKB1-WT but EGFR-mutant. Consistent with genomic analysis, cell lines with KRAS mutation are also LKB1-mutant but express WT EGFR. The status of other important tumor suppressors (p53 and PTEN) were also listed. Next, we validated the expression profiles of all the key genes as indicated in **Figure 1C** in these 6 cell lines. As shown in **Figure 1D**, H23, H460, and A549 did not express LKB1 as a result of truncation mutation, but WT EGFR could be detected in these cell lines, despite the relatively low level in H460 cells. HCC827 and H1650 expressed LKB1 and exon 19-deleted EGFR, as detected by the specific antibody against E746-A750-deleted EGFR. H1975 expresses LKB1 and L858R EGFR, which is also detected by specific antibody, however, no T790M EGFR specific antibody is currently available. Notably, the weak band of L858R EGFR in HCC827 cell line could be a non-specific signal due to the high expression level of EGFR in this cell line. Collectively, results from both NSCLC specimens and cell lines consistently indicate that EGFR and LKB1 mutations are mutually exclusive in NSCLC.

### LKB1-Mutant/EGFR-WT NSCLC Cells Are Resistant to EGFR Inhibitor Erlotinib

Given that EGFR TKIs-based targeted therapy is only applicable to NSCLC patients with EGFR mutations, we next determined the susceptibility of the 6 NSCLC cell lines to an EGFR inhibitor erlotinib. As shown in **Figure 2A**, erlotinib dramatically decreased cell viability in HCC827 cells expressing a high level of mutant EGFR (with E746-A750 deletion). However, the other two EGFR-mutant cell lines H1650 and H1975 were found to be resistant to erlotinib, most probably due to loss of PTEN and EGFR T790M mutation in these two cell types, respectively (**Figure 1C**) (Kobayashi et al., 2005; Sos et al., 2009). On the other hand, all three EGFR-WT cell lines (with LKB1 mutations) are resistant to erlotinib. Next, we further analyzed the effect of erlotinib on the signaling pathways downstream of EGFR. As shown in **Figure 2B**, erlotinib significantly inhibited the phosphorylation of EGFR, ERK1/2, and AKT in HCC827 cells bearing mutant EGFR. However, only phosphorylation of EGFR and ERK1/2, but not AKT phosphorylation was blocked by erlotinib in H1650 cells, most probably due to loss of PTEN (**Figure 2B**) (Sos et al., 2009). In contrast, erlotinib had marginal inhibitory effects on the phosphorylation status of EGFR, ERK1/2, and AKT in H1975 cells which are known to harbor T790M mutation that offers TKIs-resistance (**Figure 2B**). Among the 3 EGFR-WT cell lines (H23, H460, and A549), erlotinib has little effects on the phosphorylation of EGFR, ERK1/2 and AKT (**Figure 2B**). Finally, we tested the therapeutic efficacy of Tarceva, the clinical tablet form of erlotinib in the nude mice xenograft model. As shown in **Figure 2C**, Tarceva markedly reduced tumor volume of HCC827 cells (with mutant EGFR and WT LKB1), while its effect on H460 cells (with WT EGFR and mutant LKB1) was insignificant. In this study we used a relatively higher dose of Tarceva found in the literature (Gallagher-Colombo et al., 2015; Zhang et al., 2012) and such a dosage did not have any evident effect on mouse’s body weight (**Figure S2A**). Moreover, immunohistochemistry analysis with a cell proliferation marker Ki67 showed consistent data (**Figure S2B**). Similar with the *in vitro* results (**Figure 2B**), Tarceva totally inhibited EGFR phosphorylation and caused its degradation in the tumor formed by HCC827, whereas it only slightly decreased EGFR phosphorylation in the tumor formed by H460 (**Figure S2C**). Such data thus confirm that the EGFR TKIs are not effective for NSCLC with WT EGFR and mutant LKB1.

**Figure 2.**
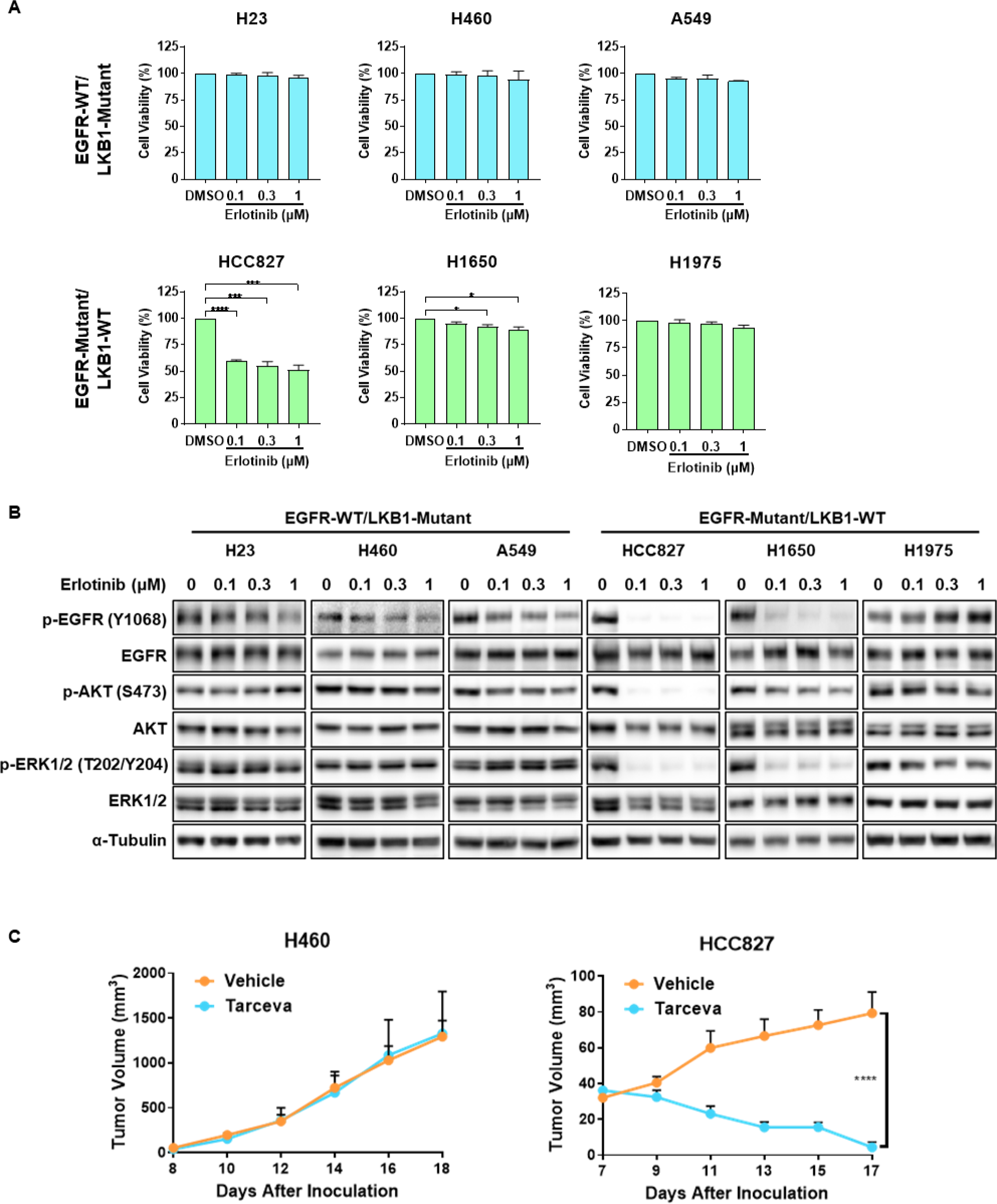
LKB1-Mutant/EGFR-WT NSCLC Cells Are Resistant to EGFR Inhibitor Erlotinib. (A) LKB1-mutant NSCLC cell lines are more resistant to EGFR inhibitor erlotinib. Six NSCLC cell lines were treated with indicated concentrations of erlotinib for 72 hrs. Cell viability was measured by CellTiter-Glo cell viability assay and normalized to the DMSO control group. Data were presented as Mean ± SE from at least 3 independent experiments. *p < 0.05, ***p < 0.001, ****p < 0.0001, (student’s t test) comparing to their respective DMSO-treated control group.
(B) Erlotinib has marginal effects on EGFR signaling pathway in LKB1-mutant NSCLC cell lines. Six NSCLC cell lines treated with indicated concentrations of erlotinib for 48 hrs and then lysed. Phosphorylation of proteins in the EGFR signaling pathway was analyzed by immunoblotting.
(C) Tarceva exhibits weaker effects on inhibiting tumor formation and cell proliferation of LKB1-mutant NSCLC cells. H460 and HCC827 cell lines were injected into the flank of BALB/c nude mice. After 7-8 days when the palpable tumor was established (50 mm^3^), mice were administered with vehicle or Tarceva (50 mg/kg/day) by oral gavage daily. Tumor volume was recorded daily. Data were presented as Mean ± SE from 6 (H460) or 8 (HCC827) mice per group. ****p < 0.0001 (two-way ANOVA). See also Figure S2.

### LKB1-Mutant/EGFR-WT NSCLC Cells Are Sensitive to Cell Death Induced by Glucose Starvation

It has been well established that under the glucose starvation, AMPK is activated by LKB1 to exert its pro-cell survival functions (Sakamoto et al., 2005; Zhang et al., 2017; Zhang et al., 2014). In addition, LKB1 deficiency sensitizes cancer cells to various types of metabolic stress (Inge et al., 2009; Shackelford et al., 2013; Whang et al., 2016). In this study, we examined the susceptibility of these 6 NSCLC cell lines to various stress conditions including glucose starvation. Interestingly, glucose starvation caused dramatic cell death in LKB1-mutant cell lines H23, H460, and A549, but not in other LKB1-WT cell lines (**Figures 3A and S3A**), suggesting that LKB1-mutant NSCLC cell lines are highly susceptible to cell death under glucose starvation. Among them, H460 was the most susceptible one, with more than 50% cell death under glucose starvation for 6 hrs (**Figure 3A**). Interestingly, deprivation of other nutrients, including glutamine, amino acids, serum, or the combination of amino acids and serum, had minimal effects on LKB1-mutant NSCLC cell lines (**Figures 3B and S3B**). Glucose replenishment rescued cell death in a dose-dependent manner in three LKB1-mutant NSCLC cell lines (**Figures 3C and S3C**). Moreover, cell death induced by glucose starvation was strongly blocked by reconstitution of WT, but not kinase-dead (KD) LKB1 in the 3 LKB1-mutanNSCLC cell lines (**Figures 3D and S3D**). In contrast, in LKB1-WT NSCLC cell lines, LKB1 knockout (KO) dramatically exacerbated cell death induced by glucose starvation (**Figures 3E and S3E**). Here we included another NSCLC cell line H358 which expresses both WT EGFR and WT LKB1. Knockout of LKB1 in H358 also sensitized cells to glucose starvation, suggesting that the sensitivity to glucose starvation is independent of EGFR status. Taken together, these results demonstrate that LKB1 plays a key role in determining the susceptibility to glucose starvation-induced cell death. We further examined the form of cell death and found that glucose starvation induces apoptotic cell death in H23 and A549 cells evidenced by (i) increased poly (ADP-ribose) polymerase (PARP) and caspase 3 cleavage, and (ii) blockage of cell death by the pan-caspase inhibitor Z-VAD (**Figures 3F and 3G**). Intriguingly, glucose starvation did not induce typical apoptotic markers such as cleavage of PARP, caspase 8 or caspase 3 in the H460 cell line (**Figure S3F**). Moreover, none of the following cell death inhibitors, including caspase inhibitor Z-VAD, necroptosis inhibitor necrostatin-1 (Nec-1), lysosome inhibitor chloroquine (CQ), ferroptosis inhibitor ferrostatin-1 (Fer-1), the receptor-interacting protein kinase 3 (RIP3) inhibitor dabrafenib (DAB), or the mixed lineage kinase domain like pseudokinase (MLKL) inhibitor necrosulfonamide (NSA), could block cell death in H460 cells under glucose starvation (**Figure S3G**). The exact nature of cell death induced by glucose starvation in H460 cells remains to be further investigated.

**Figure 3.**
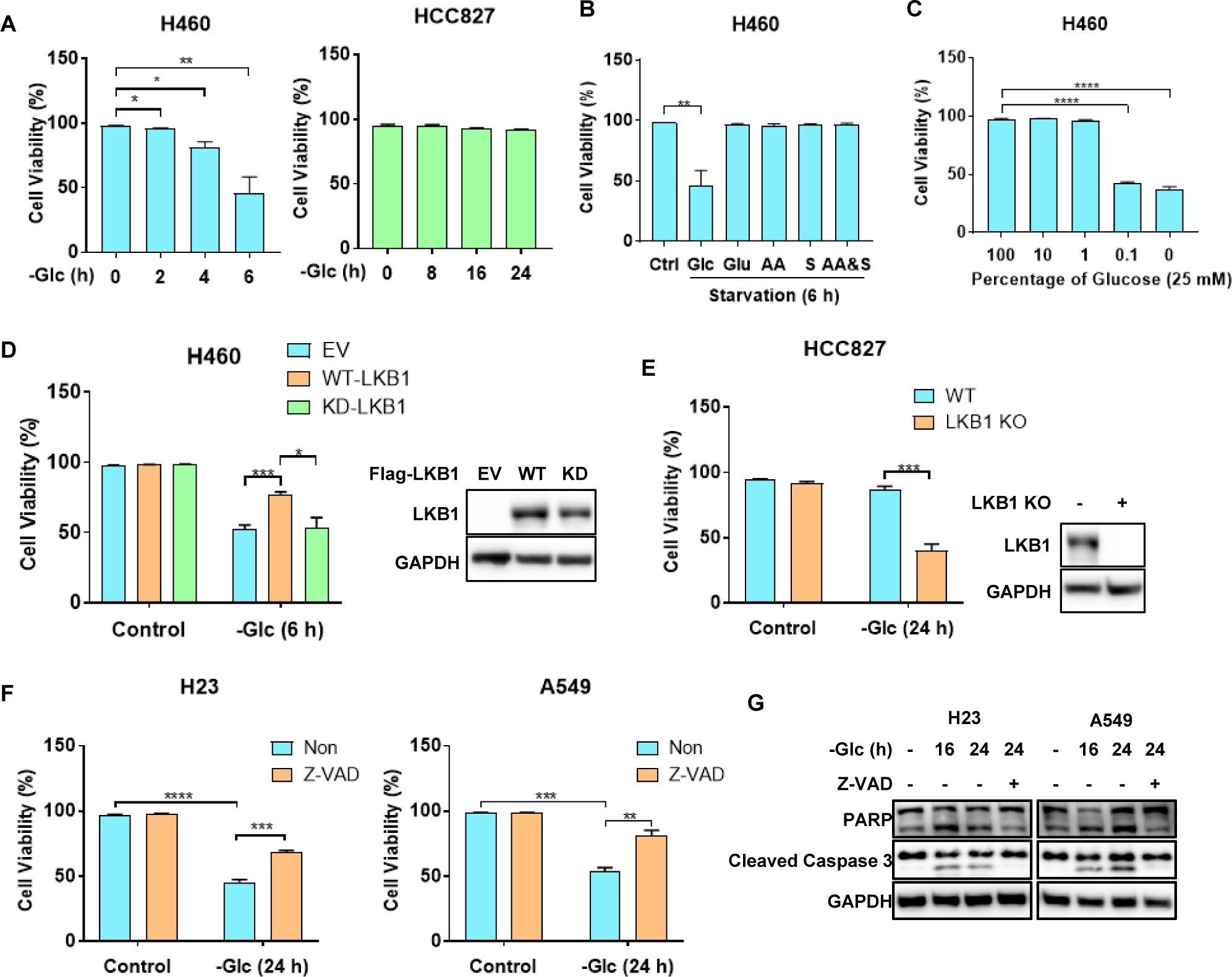
LKB1-Mutant/EGFR-WT NSCLC Cells Are Sensitive to Cell Death Induced by Glucose Starvation. (A) LKB1-mutant NSCLC cell lines are more sensitive to glucose starvation. H460 and HCC827 cell lines were treated with glucose starvation for the indicated times. Cell viability was measured by PI exclusion assay using flow cytometry.
(B) LKB1-mutant NSCLC cell lines are only sensitive to glucose starvation. H460 cell line was cultured in DMEM without glucose, glutamine, amino acids (AA), serum, or the combination of AA and serum for 6 hrs. Cell viability was measured by PI exclusion assay using flow cytometry.
(C) Glucose replenishment rescues cell death in a dose-dependent manner in LKB1-mutant NSCLC cell lines. H460 cell line was cultured in glucose-free DMEM, supplemented with the indicated percentage of glucose for 6 hrs. Cell viability was measured by PI exclusion assay using flow cytometry.
(D) LKB1 reconstruction activates AMPK and rescues cell death in LKB1-mutant NSCLC cell lines upon glucose starvation. H460 cell line stably expressing empty vector (EV), wild type (WT)- or kinase dead (KD)-LKB1 was treated with full or glucose-free DMEM for 6 hrs. Cell viability was measured by PI exclusion assay using flow cytometry. LKB1 reconstitution was shown.
(E) LKB1 knockout sensitizes LKB1-WT NSCLC cell lines to cell death induced by glucose starvation. Wild type (WT) or LKB1 knockout (KO) HCC827 cell line was treated with full or glucose-free DMEM for 24 hrs. Cell viability was measured by PI exclusion assay using flow cytometry. LKB1 knockout was shown.
(F and G) Glucose starvation induces apoptosis in LKB1-mutant NSCLC cell lines. H23 and A549 cell lines cultured in full or glucose-free DMEM were treated with Z-VAD (40 μM) for 24 hrs. **(F)** Cell viability was measured by PI exclusion assay using flow cytometry. **(G)** Cells were lyzed and cleavage of PARP and caspase 3 were analyzed by immunoblotting. Data were presented as Mean ± SE from at least 3 independent experiments. *p < 0.05, **p < 0.01, ***p < 0.001, ****p < 0.0001 (student’s t test) comparing to their respective control (full medium or non-treated) group. See also Figure S3.

### Glucose Starvation Induces Cell Death via Oxidative Stress in EGFR-WT/LKB1-Mutant NSCLC Cells

It has been well established that glucose starvation enhances ROS generation (Chen et al., 2009; Graham et al., 2012; Liu et al., 2003). The activation of AMPK by ROS has also been widely reported (Emerling et al., 2009; Mackenzie et al., 2013; Rabinovitch et al., 2017; Ren and Shen, 2019; Zmijewski et al., 2010). However, it remains elusive whether redox activation of AMPK is dependent on or independent of LKB1. Therefore, we examined the possibility that the high sensitivity of LKB1-mutant cells to glucose starvation is mediated via oxidative stress. Glucose starvation enhanced ROS production, and the antioxidant NAC markedly reduced the ROS levels in cells under glucose starvation (**Figures 4A and S4A**). In contrast, glucose starvation did not increase ROS level in LKB1-WT NSCLC cells (**Figures 4B and S4B**), suggesting that in the presence of LKB1, cells are able to maintain the intracellular redox homeostasis, probably through AMPK. Consistently, NAC blocked glucose starvation-induced cell death (**Figures 4C and S4C**), indicating that oxidative stress induced by glucose starvation is the major mediator of cell death.

**Figure 4.**
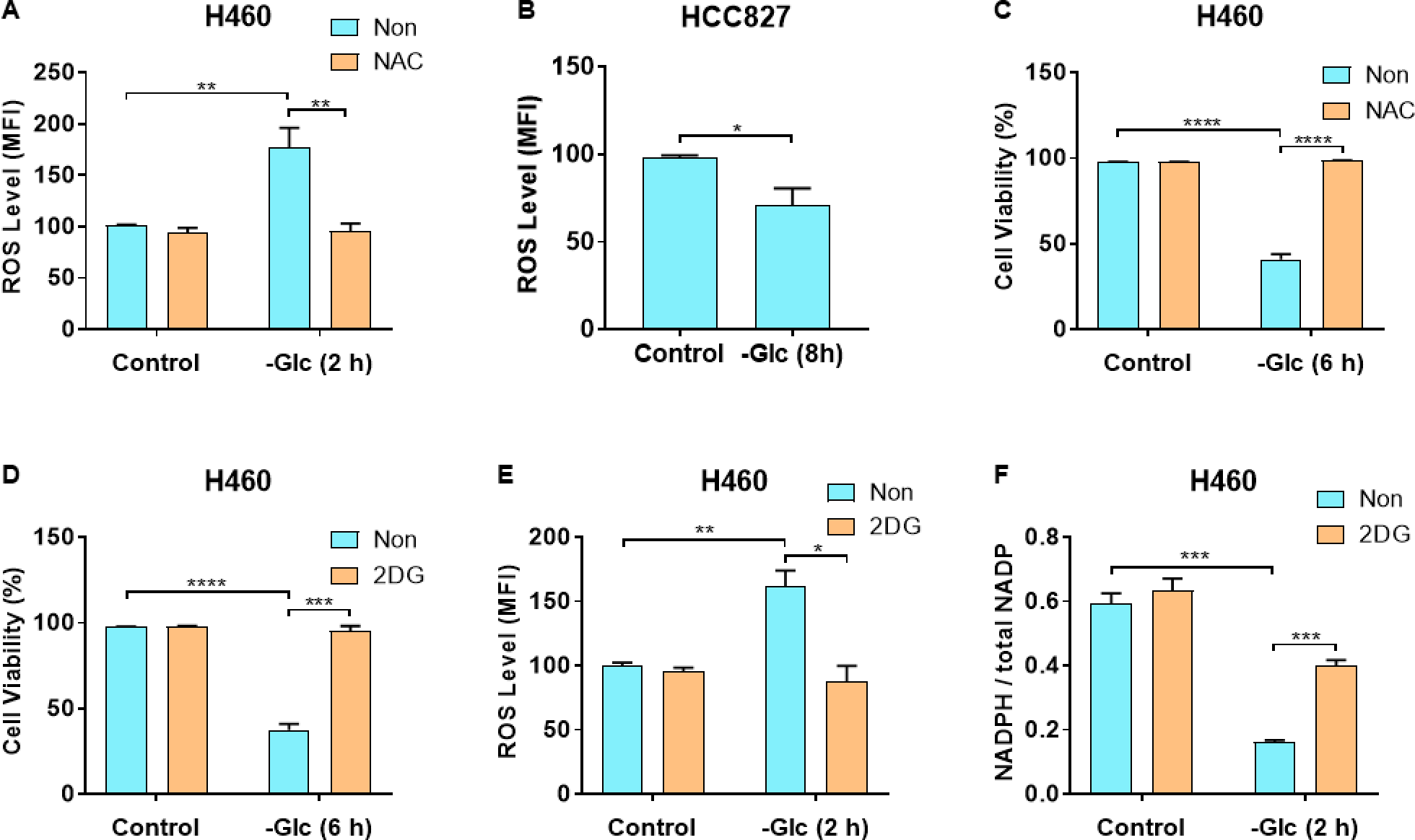
Glucose Starvation Induces Cell Death via Oxidative Stress in EGFR-WT/LKB1-Mutant NSCLC Cells. (A and B) Glucose starvation increases the ROS level in LKB1-mutant NSCLC cells. **(A)** H460 cell line cultured in full or glucose-free DMEM were treated with NAC (5 mM) for 2 hrs. **(B)** HCC827 cell line was treated with full or glucose-free DMEM for 8 hrs. Intracellular ROS levels were measured using CellROX Green reagent by flow cytometry. A representative histogram of flow cytometry results was shown.
(C) NAC blocks glucose starvation-induced cell death in LKB1-mutant NSCLC cell lines. H460 cell line cultured in full or glucose-free DMEM were treated with NAC (5 mM) for 6 hrs. Cell viability was measured by PI exclusion assay using flow cytometry.
(D) 2DG rescues glucose starvation-induced cell death LKB1-mutant NSCLC cell lines. H460 cell line cultured in full or glucose-free DMEM were treated with 2DG (5 mM) for 6 hrs. Cell viability was measured by PI exclusion assay using flow cytometry.
(E) 2DG reduces ROS level in LKB1-mutant NSCLC cell lines under glucose starvation. H460 cell line cultured in full or glucose-free DMEM were treated with 2DG (5 mM) for 2 hrs. Intracellular ROS levels were measured using CellROX Green reagent by flow cytometry.
(F) 2DG maintains NAPDH level in LKB1-mutant NSCLC cell lines under glucose starvation. H460 cell line cultured in full or glucose-free DMEM were treated with 2DG (5 mM) for 2 hrs. NADPH and NADP levels were measured using NADP/NADPH Quantitation Kit following the manufacturer’s instruction. Data were presented as Mean ± SE from at least 3 independent experiments. *p < 0.05, **p < 0.01, ***p < 0.001, ****p < 0.0001 (student’s t test) comparing to their respective control (full medium or non-treated) group. See also Figure S4.

At present, the non-metabolizable glucose analog 2-deoxy-D-glucose (2DG) has been widely used as a mimetic of glucose starvation and as an anti-cancer therapeutic (Aft et al., 2002; Cheong et al., 2011; Oladghaffari et al., 2015). On the other hand, it has reported that, although 2DG cannot be utilized in the glycolysis, it can enter the pentose phosphate pathway (PPP) to maintain the production of a key antioxidant nicotinamide adenine dinucleotide phosphate reduced (NADPH), thus blocking glucose starvation-induced ROS generation and cell death (Jeon et al., 2012; Li et al., 2018). In this study, while 2DG alone did not cause any cell death in LKB1-mutant cells lines, it almost totally blocked cell death in H460 (**Figures 4D**) and A549 cells, but not in H23 cells (**Figure S4D**). As expected 2DG reduced ROS level and recovered NADPH level in H460 cells (**Figures 4E and 4F**), as well as in A549 cells (**Figure S4D and S4E**). Collectively, our data suggest that the oxidative stress and ROS are the main mediators of cell death in EGFR-WT/LKB1-mutant NSCLC cells under glucose starvation.

### Impaired AMPK Activation in EGFR-WT/LKB1-Mutant NSCLC Cells Is Implicated in The Susceptibility to Cell Death under Glucose Starvation

Based on the well-established fact that AMPK is a key downstream target of LKB1 (Hardie and Alessi, 2013), here we examined the implication of AMPK in cell death induced by glucose starvation in LKB1-WT and -mutant NSCLC cell lines. To our surprise, in all 3 EGFR-WT/LKB1-mutant NSCLC cell lines, glucose starvation markedly reduced the p-AMPK and p-ACC level (**Figures 5A and S5A**). In contrast, all the EGFR-mutant/LKB1-WT cell lines showed increased p-AMPK and p-ACC level in response to glucose starvation (**Figures 5A and S5A**). Consistent with the cell death results (**Figures 3 and S3**), reconstitution of WT but not KD LKB1 into LKB1-mutant NSCLC cells was able to increase p-AMPK and p-ACC level under both normal and glucose starvation conditions (**Figures 5B and S5B**). On the contrary, LKB1 deletion in LKB1-WT cells impaired AMPK activation in response to glucose starvation in (**Figures 5C and S5C**), supporting the well-established role of LKB1 in AMPK activation upon glucose starvation.

**Figure 5.**
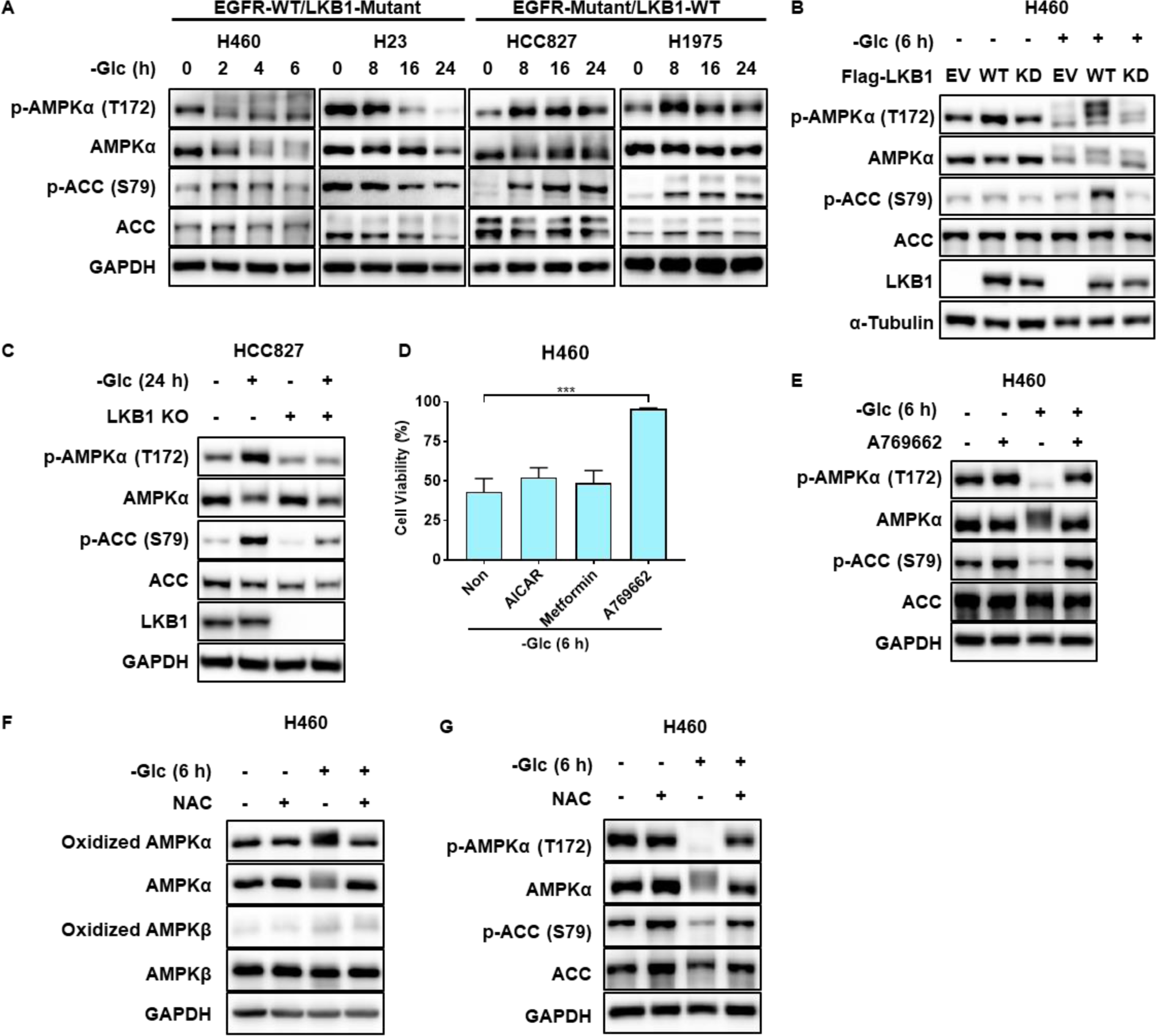
Impaired AMPK Activation in EGFR-WT/LKB1-Mutant NSCLC Cells Is Implicated in The Susceptibility to Cell Death under Glucose Starvation. (A) AMPK is inactivated in LKB1-mutant NSCLC cell lines upon glucose starvation. NSCLC cell lines were treated with glucose-free DMEM for the indicated times and then lysed. AMPK activity was measured by ACC and AMPKα phosphorylation using immunoblotting.
(B) LKB1 reconstruction activates AMPK in LKB1-mutant NSCLC cell lines upon glucose starvation. H460 cell line stably expressing empty vector (EV), wild type (WT)- or kinase dead (KD)-LKB1 was treated with full or glucose-free DMEM for 6 hrs. AMPK activity was measured by ACC and AMPKα phosphorylation using immunoblotting.
(C) LKB1 knockout impairs AMPK activation and sensitizes LKB1-WT NSCLC cell lines to cell death induced by glucose starvation. Wild type (WT) or LKB1 knockout (KO) HCC827 cell line was treated with full or glucose-free DMEM for 24 hrs. AMPK activity was measured by ACC and AMPKα phosphorylation using immunoblotting.
(D) Only AMPK allosteric activator A769662 can block cell death in LKB1-mutant NSCLC cell lines upon glucose starvation. H460 cell line cultured in glucose-free DMEM was treated with AICAR (500 μM), metformin (5 mM) or A769662 (200 μM) for 6 hrs. Cell viability was measured by PI exclusion assay using flow cytometry. Data were presented as Mean ± SE from at least 3 independent experiments. ***p < 0.001 (student’s t test) comparing to non-treated group.
(E) AMPK allosteric activator A769662 activates AMPK in LKB1-mutant NSCLC cell lines upon glucose starvation. H460 cell line cultured in full or glucose-free DMEM was treated with A769662 (200 μM) for 6 hrs. AMPK activity was measured by ACC and AMPKα phosphorylation using immunoblotting.
(F) AMPKα is oxidized in LKB1-mutant NSCLC cell lines upon glucose starvation. H460 cell line cultured in full or glucose-free DMEM was treated with NAC (5 mM) for 6 hrs. Cysteine oxidation of AMPK subunits was detected.
(G) NAC maintains AMPK activity in LKB1-mutant NSCLC cell lines upon glucose starvation. H460 cell line as treated in **(F)** was lysed and AMPK activity was measured by ACC and AMPKα phosphorylation using immunoblotting. See also Figure S5.

Given that AMPK activity is necessary to maintain cell survival under glucose starvation condition (Jeon et al., 2012), we tested different types of AMPK activators in LKB1-mutant cell lines. Among them, metformin and AICAR are known to activate AMPK via mimicking AMP or inhibiting mitochondrial complex I, respectively, which are LKB1-dependent (Hawley et al., 2003; Shaw et al., 2005). As expected, neither metformin nor AICAR was able to block cell death under glucose starvation (**Figure 5D**). In contrast, the LKB1-independent AMPK allosteric activator A769662 effectively protected cells against glucose starvation-induced cell death in all three LKB1-mutant NSCLC cell lines (**Figures 5D and S5D**). Consistently, A769662 maintained AMPK activity in these cell lines (**Figures 5E and S5E**), suggesting that the allosteric activation of AMPK in LKB1-mutant cells is able to offer strong protection against cell death induced by glucose starvation, and further illustrating the importance of AMPK in determining the susceptibility of NSCLC to cell death under glucose starvation. To shed more lights on the mechanism of how ROS regulate AMPK in the absence of LKB1, we focused on AMPK post-translational modifications (PTMs). In this study, we observed an obvious upshift of AMPKα band in LKB1-mutant NSCLC cell lines upon glucose starvation (**Figure 5A**), suggesting that AMPKα undergo certain PTMs. Previous studies have shown that increased ROS can inhibit protein functions through direct cysteine oxidation (Anastasiou et al., 2011; Li et al., 2018). We therefore analyzed whether AMPKα undergoes oxidation upon glucose starvation. We found that oxidation of AMPKα, but not that of AMPKβ, was increased by glucose starvation, and this effect was abolished by treatment with NAC (**Figures 5F and S5F**). Interestingly, NAC also maintained AMPK activity upon glucose starvation (**Figures 5G and S5G**), indicating that glucose starvation may negatively regulate AMPK activity via oxidation in LKB1-mutant cells. These data suggest that the oxidation of AMPKα decreases its activity, which diminishes AMPK activity and impairs the protective function of AMPK in cell survival under glucose starvation. Taken together, these results indicate that the molecular mechanisms underlying the defective AMPK activation in EGFR-WT/LKB1-mutant NSCLC cells were two folds: first, lack of LKB1-mediated T172 phosphorylation and second, ROS-mediated AMPKα oxidation.

### SGLT2 Inhibitor Canagliflozin Causes Synthetic Lethality in LKB1-Mutant NSCLC Cells

To mimic the effect of glucose starvation *in vivo*, we next investigated the possibility of targeting glucose transporter. As mentioned above, the SGLT family proteins are involved in the active uptake of glucose, with predominant expression in intestine and kidney (Vrhovac et al., 2015). Moreover, it has been reported that functional SGLT2 expression is detected in various cancer types, including lung, pancreatic, prostate, colon, and liver cancers (Ishikawa et al., 2001; Kaji et al., 2018; Saito et al., 2015; Scafoglio et al., 2015; Scafoglio et al., 2018; Villani et al., 2016). Indeed, all 6 NSCLC cell lines expressed SGLT2 (**Figure 6A**). Targeting SGLT2 by its inhibitors has been shown to inhibit tumor cell growth in pancreatic, prostate, colon, liver and lung cancers (Kaji et al., 2018; Saito et al., 2015; Scafoglio et al., 2015; Scafoglio et al., 2018; Villani et al., 2016). Thus, it is of interest and importance to investigating whether SGLT2 inhibitors would mimic the effect of glucose starvation in selective killing of the EGFR-WT/LKB1-mutant NSCLC cells. Here we tested three FDA-approved SGLT2 inhibitors, *i.e.*, canagliflozin, dapagliflozin, and empagliflozin individually in the most sensitive cell line H460 (**Figure S6A**). Among them, only canagliflozin induced significant cell death in cells cultured under low glucose concentration (1 g/L, 5.56 mM), which is within the range of physiological blood glucose concentration (4.4 to 6.1 mM) (Engelgau et al., 2000). Unlike canagliflozin, the other two inhibitors (dapagliflozin and empagliflozin) did not induce cell death at the same concentration as canagliflozin (**Figure S6A**). Then we determined the susceptibility of all 6 NSCLC cell lines to canagliflozin. Consistent with glucose starvation, canagliflozin selectively caused cell death in EGFR-WT/LKB1-mutant NSCLC cell lines in a dose-dependent manner, whereas EGFR-mutant/LKB1-WT cell lines HCC827, H1650 and H1975 were much more resistant to canagliflozin (**Figures 6B and S6B**). Moreover, the glucose uptake in EGFR-WT/LKB1-mutant NSCLC cell lines, measured using a non-metabolizable fluorescent glucose analog 2-(N-(7-Nitrobenz-2-oxa-1,3-diazol-4-yl)Amino)-2-deoxyglucose (2NBDG), was significantly reduced by canagliflozin, but not by dapagliflozin or empagliflozin (**Figure 6C**), which may explain the inability of dapagliflozin and empagliflozin in cell killing shown earlier (**Figure S6A**). To understand the underlying mechanisms, we examined the implication of AMPK in cells treated with canagliflozin. Canagliflozin led to decreased AMPK activity in LKB1-mutant cell lines, while activated AMPK in LKB1-WT cell lines, a similar pattern as glucose starvation (**Figures 6D and S6C**). Finally, we tested the anti-tumor efficacy of canagliflozin *in vivo*. Invokana is the clinical tablet form of canagliflozin which is currently used to treat type 2 diabetes. In a BALB/c nude mice xenograft model, daily oral gavage of Invokana (100 mg/kg/day), a dosage reported by others in study of its anti-tumor activity (Friess et al., 2005; Kaji et al., 2018; Villani et al., 2016; Zhang et al., 2012), significantly mitigated tumor growth of H460 and A549 cell lines, but not that of HCC827 (**Figures 6E and S6D**). Consistent with reduced tumor growth, in Invokana-treated H460 and A549 xenograft tumors, the proliferation marker Ki67 was much weaker, whereas no obvious difference was detected in HCC827 xenograft tumor (**Figures 6F and S6E**). Moreover, in line with the *in vitro* data, significant AMPK activation by Invokana was only observed in HCC827 xenograft tumor, but not in LKB1-mutant H460 and A549 cells (**Figures 6G and S6F**). Notably, Invokana had a minimal physiological side effect on mice, demonstrated by the similar trends in body weight as the vehicle (**Figure S6G**). The results shown above demonstrate that Invokana is effective on inhibiting tumor growth of H460 and A549 cell lines that are resistant to EGFR TKIs. Collectively, such data support our notion that in EGFR-WT/LKB1-mutant NSCLC cells, targeting SGLT2 with canagliflozin results in synthetic lethality, suggesting that canagliflozin can be developed as a novel targeted therapy for NSCLC with LKB1 mutations.

**Figure 6.**
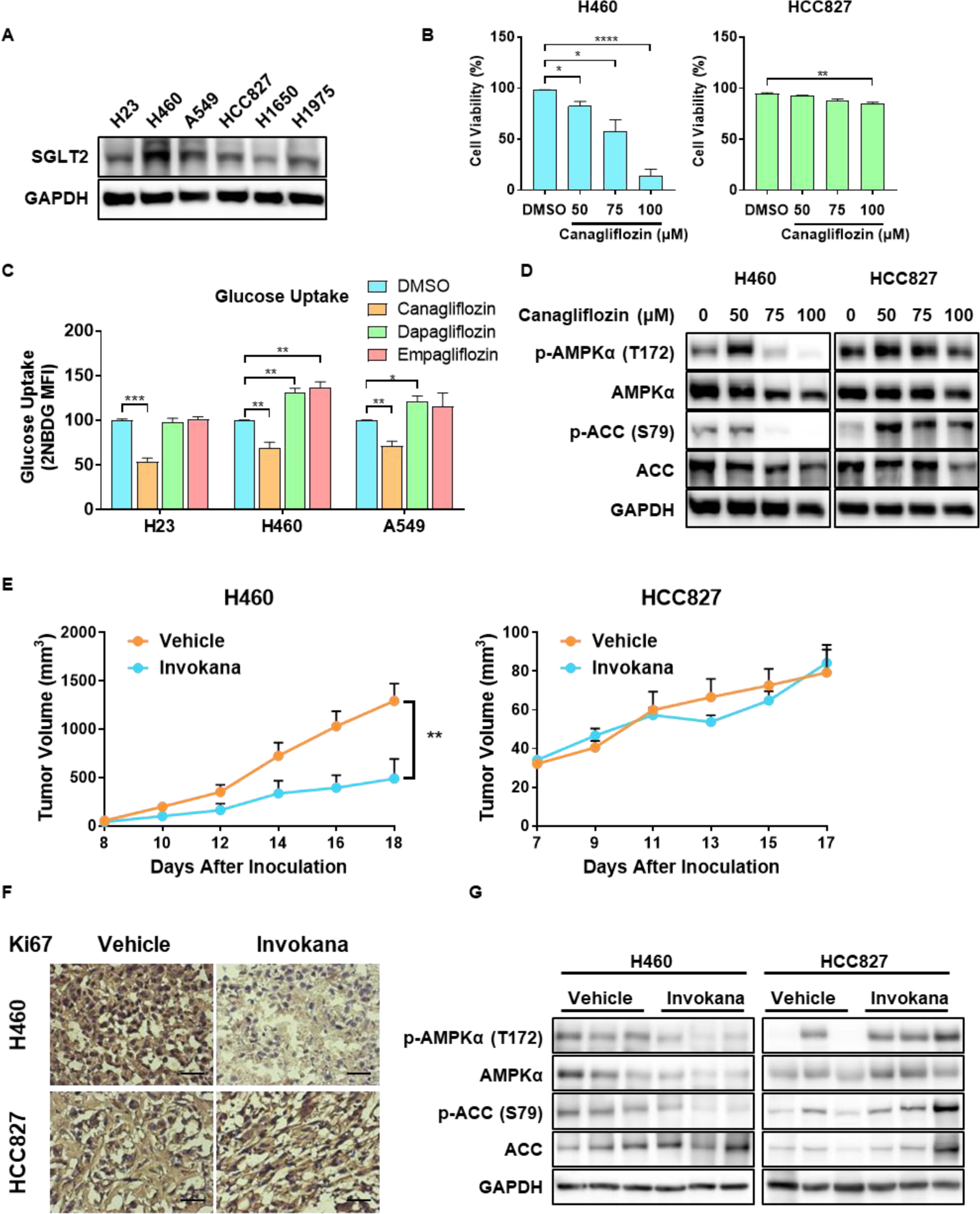
SGLT2 Inhibitor Canagliflozin Causes Synthetic Lethality in LKB1-Mutant NSCLC Cells. (A) SGLT2 can be detected in NSCLC cell lines. Same samples in Figure 1D were also detected using an SGLT2 antibody.
(B) LKB1-mutant NSCLC cell lines are sensitive to canagliflozin. H460 and HCC827 cell lines cultured in low glucose DMEM (1 g/L) were treated with indicated concentrations of canagliflozin for 24 hrs. Cell viability was measured by PI exclusion assay using flow cytometry.
(C) Canagliflozin, but not dapagliflozin or empagliflozin inhibits glucose uptake. H23, H460, and A549 cell lines cultured in glucose-free DMEM were treated with canagliflozin (100 μM) for 2 hrs. 2NBDG (250 μM) was added 1 hr prior to harvesting cells. Glucose uptake was measured by the intracellular fluorescence intensity of 2NBDG using flow cytometry.
(D) AMPK is inactivated in LKB1-mutant NSCLC cell lines under canagliflozin treatment. H460 and HCC827 cell lines as treated in **(B)** were lysed. AMPK activity was measured by ACC and AMPKα phosphorylation using immunoblotting.
(E and F) Invokana significantly mitigates tumor growth and cell proliferation of LKB1-mutant NSCLC cells *in vivo*. H460 and HCC827 cell lines were injected into the flank of BALB/c nude mice. After 7-8 days when the palpable tumor was established (50 mm^3^), mice were administered with vehicle or Invokana (100 mg/kg/day) by oral gavage daily.
(G) Tumor volume was recorded daily. Tumors were excised from mice 24 hrs after the last gavage. Samples were embedded, frozen, sectioned and stained for cell proliferation marker Ki67. **(F)** Representative images are shown. Scale bar, 100 µm. Data were presented as Mean ± SE from at least 6 mice per group. ns, not significant; *p < 0.05 (two-way ANOVA).
(G) Invokana only activates AMPK in tumors formed by LKB1-WT NSCLC cells. Tumors formed by H460 and HCC827 cell lines were excised from mice 24 hrs after the last gavage. Samples were lysed and AMPK activity was measured by ACC and AMPKα phosphorylation using immunoblotting. See also Figure S6.

## DISCUSSION

### Mutual Exclusivity between EGFR and LKB1 Mutations in NSCLC

Gain-of-function mutation of EGFR is a key oncogenic mechanism in the development of NSCLC. Currently, specific EGFR TKIs have been successfully developed as a targeted therapy for those NSCLC patients with EGFR mutations (in 10-15% of Caucasian NSCLC patients, and in 30-50% Asian patients) (Han et al., 2005; Janne et al., 2005; Kosaka et al., 2004; Tsao et al., 2005). However, for the majority of the NSCLC patients with WT EGFR, currently there is no targeted therapy available. On the other hand, LKB1 is an important tumor suppressor and its loss-of-function mutation is frequently detected in 15-30% of NSCLC patients (Ding et al., 2008; Gill et al., 2011; Ji et al., 2007). The oncogenic EGFR and KRAS mutations have been widely reported to be mutually exclusive in NSCLC and other cancer types (Choughule et al., 2014; Fu et al., 2014; Kris et al., 2011; Shigematsu et al., 2005; Tam et al., 2006). Moreover, KRAS mutation often predicts negative responsiveness of NSCLC to EGFR TKIs, thus regarded as a negative biomarker for EGFR TKIs-based targeted therapies for NSCLC (Li et al., 2014b; Mao et al., 2010; Pao et al., 2005b). Interestingly, LKB1 mutation alone could not induce pulmonary neoplasia, but the combination with KRAS mutation exacerbated tumor burden and caused more frequent differentiation and metastasis (Ji et al., 2007). Moreover, genomic analyses of NSCLC tumor samples have revealed that LKB1 mutations co-occur with KRAS mutations in about 10% of NSCLC patients (Ding et al., 2008; Ji et al., 2007; Makowski and Hayes, 2008). These findings thus raised the possibility that EGFR and LKB1 mutation are also mutually exclusive in NSCLC. To verify this possibility, we performed genomic analysis of NSCLC samples as well as detected EGFR and LKB1 expression in established NSCLC cell lines and found a significant mutual exclusivity between EGFR and LKB1 mutations in NSCLC (**Figure 1**). Notably, the mutual exclusivity between EGFR and LKB1 mutations was also found in a recent report in lung cancer patients in Singapore, although this study was not included by cBioPortal (Heong et al., 2018). Consistently, we found that LKB1-mutant NSCLC cell lines (with WT EGFR) are resistant to EGFR inhibitor erlotinib (**Figure 2**). Up to date, various attempts have been made to target NSCLC patients with WT EGFR, including using the combination of EGFR TKIs with other inhibitors such as HER2 inhibitor pertuzumab, vascular endothelial cell growth factor (VEGF) inhibitor bevacizumab, AKT inhibitor MK2206 or autophagy inhibitor hydroxychloroquine (HCQ) (Chen et al., 2013; Goldberg et al., 2012; Huang et al., 2016; Lara et al., 2015; Li et al., 2014a; Wang et al., 2008). However, these combinational strategies only offer limited effectiveness and only benefit a small number of NSCLC patients. As such, there is an urgent need and of great significance to develop novel targeted therapies for those NSCLC patients with WT EGFR. The mutual exclusivity of EGFR and LKB1 mutation found in our study thus offers a window of opportunity for such an approach.

### Defective AMPK Activation Is Implicated in The Selective Killing of LKB1-Mutant NSCLC Cells under Glucose Starvation

LKB1 was originally identified as the tumor suppressor responsible for the inherited cancer disorder Peutz-Jeghers Syndrome (PJS) (Hemminki et al., 1998). Since then, LKB1 mutation has been observed in many types of cancers, including lung cancer, ovarian cancer, breast cancer, cervical cancer and pancreatic cancer (Bardeesy et al., 2002; Ji et al., 2007; Wingo et al., 2009). Supporting the tumor suppressor function of LKB1, mice with hetero- or homozygous deletion of LKB1 are prone to develop lung, cervical, and pancreatic tumors (Hezel et al., 2008; Ji et al., 2007; Morton et al., 2010; Wingo et al., 2009). On the other hand, numerous studies have reported that LKB1 mutation confers sensitivity of cancer cells to various stress conditions and cancer therapeutics, such as rapamycin and phenformin (Contreras et al., 2010; Shackelford et al., 2013). A key finding of our study is that LKB1-mutant NSCLC cells are more susceptible to glucose starvation, compared to the LKB1-WT cells (**Figure 3**).

To understand the molecular mechanisms underlying the hyper-sensitivity of LKB1-mutant NSCLC cells to glucose starvation, we focused on AMPK, the most important downstream target of LKB1. Previous studies have reported that glucose starvation enhances ROS generation and induces oxidative stress, probably due to downregulated PPP that generates NADPH, a key intracellular antioxidant (Chen et al., 2009; Graham et al., 2012; Jeon et al., 2012; Liu et al., 2003). However, other studies challenged this view by reporting that high glucose also stimulates ROS generation (Yu et al., 2011; Zhang et al., 2007). This argument is reasonable because ROS are normally generated by the electron leakage from the mitochondrial electron transport chain during the oxidative phosphorylation process (Murphy, 2009). A high level of glucose means high workload for mitochondria, leading to higher levels of ROS production. Moreover, the effect of ROS and oxidative stress on AMPK seems also controversial. On one hand, it has been reported that ROS are able to activate AMPK through different mechanisms, including calcium and CaMKKβ-dependent pathway, direct cysteine oxidation, and decrease in ATP level (Choi et al., 2001; Emerling et al., 2009; Mackenzie et al., 2013; Mungai et al., 2011; Rabinovitch et al., 2017; Zmijewski et al., 2010). On the other hand, there are several lines of evidence indicating that ROS play a negative role in regulating AMPK activity by direct oxidation of cysteine residues on AMPKα subunit or through (protein kinase C) PKC activation (Saberi et al., 2008; Shao et al., 2014). In this study, we provided clear evidence suggesting that in LKB1-mutant NSCLC cells, glucose starvation induces cell death via impaired AMPK activation and oxidative stress, based on the following observations: (i) ROS level is dramatically elevated in LKB1-mutant NSCLC cells under glucose starvation (**Figures 4A and S4A**); (ii) Antioxidant NAC is able to protect against cell death (**Figures 4C and S4C**) and restore AMPK activation (**Figures 5G and S5G**); (iii) Reconstitution of LKB1 in LKB1-mutant cells offers significant protection against cell death induced by glucose starvation, whereas deletion of LKB1 in LKB1-WT cells greatly sensitizes cell death (**Figures 3 and 5**). These findings are consistent with two studies showing that fatty acid oxidation, either supported by AMPK or by Nur77, is critically important to maintain redox homeostasis in cells under metabolic stress (Jeon et al., 2012; Li et al., 2018). Interestingly, it has also been reported that Nur77 negatively regulates AMPK activity through sequestering LKB1 in the nucleus, suggesting that a sophisticated regulatory network may exist to link Nur77 and AMPK with fatty acid oxidation (Zhan et al., 2012). Moreover, our notion linking oxidative stress-AMPK inactivation-cell death under glucose starvation in LKB1-mutant cells was supported by another key finding: allosteric activation of AMPK by A769662 protects against cell death induced by glucose starvation in LKB1-mutant cells (**Figures 5D and S5D**). Meanwhile, it is intriguing that elevated ROS level was only found in LKB1-mutant cells, but not in LKB1-WT cells (**Figures 4A and S4A**), indicating the possibility of the LKB1 is involved as an upstream mechanism in ROS production, thus emphasizing the importance of the LKB1-AMPK axis in maintaining redox homeostasis. The intricate relationship among LKB1, AMPK and ROS under glucose starvation remains to be further elucidated.

To shed more lights on the mechanism of how ROS impair AMPK activity under glucose starvation in LKB1-mutant NSCLC cell, we investigated the protein oxidation status of AMPK. Previous studies have reported that H_2_O_2_-induced direct *S*-glutathionylation of cysteine residues Cys299 and Cys304 on AMPKα subunit activates AMPK (Zmijewski et al., 2010), although this point of view has been challenged recently (Hinchy et al., 2018). On the contrary, several lines of evidence have also suggested that ROS may inhibit the LKB-AMPK axis (Dolinsky et al., 2009; Saberi et al., 2008; Wagner et al., 2006). Under glucose starvation or exogenous H_2_O_2_ treatment, AMPK is negatively regulated by oxidation of cysteine residues Cys130 and Cys174 on its α subunit, which causes oxidative aggregation and interferes the interaction between LKB1 and AMPK (Shao et al., 2014). As seen in **Figure 5**, there is an obvious upshift of the AMPKα band under starvation conditions, which we have proved that it resulted from oxidation. However, the total AMPKα level is also reduced, suggesting that AMPK protein stability may be decreased as a result of oxidation. Indeed, there are reports describing AMPK oxidation or ubiquitination that negatively regulates AMPK activity (Pineda et al., 2015; Qi et al., 2008; Shao et al., 2014; Vila et al., 2017). Therefore, data from this part of our study link impaired AMPK activation, oxidative stress and cell death in response to glucose starvation in LKB1-mutant cells.

### Development of SGLT2 Inhibitors as A Targeted Therapy for EGFR-WT/LKB1-Mutant NSCLC

Loss of tumor suppressor LKB1 promotes metabolic reprogramming and increases the growth and proliferation of cancer cells (Faubert et al., 2014; Faubert et al., 2015). However, at the same time, impaired AMPK activation by LKB1 confers cancer cells vulnerability to metabolic and oxidative stresses, raising the possibility of targeting cancer metabolism for synthetic lethality in LKB1-mutant cancers. Data from the earlier parts of our study reveal that EGFR-WT/LKB1-mutant NSCLC cells are highly susceptible to cell death induced by glucose starvation. Here we attempted to mimic glucose starvation both *in vitro* and *in vivo* by inhibition of glucose uptake and discovered that the SGLT2 inhibitor canagliflozin is highly effective in killing EGFR-WT/LKB1-mutant NSCLC cells. Although canagliflozin has been shown to inhibit tumor growth in multiple cancers including pancreatic, prostate, colon, liver and lung cancers (Kaji et al., 2018; Scafoglio et al., 2015; Scafoglio et al., 2018; Villani et al., 2016), this therapeutic agent has not been studied as a target therapy. In this study, we tested the efficacy of canagliflozin on inhibiting tumor growth both *in vitro* and *in vivo*. Our data suggest that canagliflozin and its clinical tablet form Invokana, are able to selectively kill LKB1-mutant NSCLC cells *in vitro* and reduce tumor growth in xenograft nude mice *in vivo* (**Figures 6 and S6**). Interestingly, the other two SGLT2 inhibitors, *i.e.*, dapagliflozin and empagliflozin failed to lower glucose uptake and to cause cell death (**Figure 6C and S6A**). These results are consistent with previous findings that only canagliflozin, but not dapagliflozin or empagliflozin, is able to inhibit mitochondrial complex I, in addition to SGLT2 inhibition (Hawley et al., 2016; Secker et al., 2018; Villani et al., 2016). Also, it has been reported that in HEK293 cells only canagliflozin is capable of inhibiting glucose uptake (Hawley et al., 2016). Therefore, among the approved SGLT2 inhibitors, canagliflozin would be a better choice for development as a novel targeted therapy applicable for EGFR-WT/LKB1-mutant NSCLC. Collectively, our findings suggest that SGLT2 inhibitor canagliflozin is able to effectively and specifically cause synthetic lethality in LKB1-mutant NSCLC cells, which cannot be killed by EGFR inhibitors, thus providing new insights into the personalized therapy for NSCLC.

Taken together, data from our study suggest that the LKB1 status decides the response to glucose starvation and SGLT inhibition and synthetic lethality can be achieved in LKB1-mutant/EGFR-WT NSCLC with glucose starvation or SGLT2 inhibition. As summarized in **Figure 7**, LKB1-WT (also EGFR-mutant) NSCLC cells are insensitive to glucose starvation or SGLT2 inhibition due to LKB1-mediated AMPK activation and maintenance of the redox balance. In contrast, in LKB1-mutant (also EGFR-WT) NSCLC cells, glucose starvation or SGLT2 inhibition fails to activate AMPK and selectively kills this group of cancer cells via oxidative stress (**Figure 7**). Translationally, our findings provide clear evidence supporting the development of a novel targeted therapy: using SGLT2 inhibitor canagliflozin for a significant proportion of NSCLC patients (15% – 30%) with WT EGFR and mutant LKB1, thus filling a large gap in the current clinical practice for treatment of NSCLC, one of the most prevalent and deadly cancer types worldwide.

**Figure 7.**
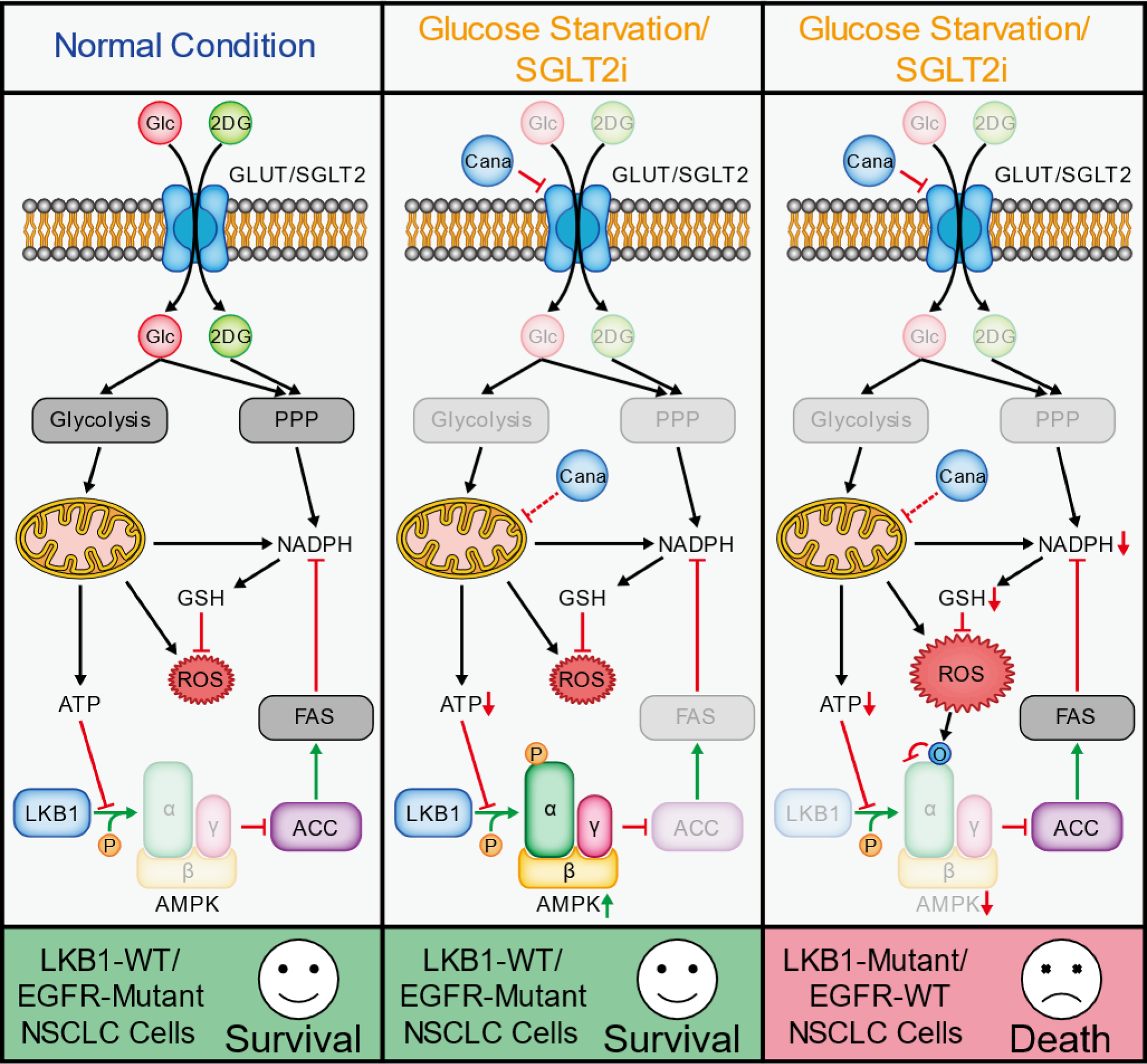
Synthetic lethality in LKB1-mutant/EGFR-WT NSCLC cells by glucose starvation or SGLT2 inhibition. A model proposing the mechanism by which glucose starvation or SGLT2 inhibition exerts synthetic lethality on LKB1-mutant NSCLC cells is shown. Black arrows show the direction of biological processes. Green arrows denote positive regulation. Red arrows or blunt ends denote negative regulation. Inhibited pathways or proteins are faded. P, phosphorylation; O, oxidation.

## ACKNOWLEDGMENTS

We gratefully acknowledge Dr. D. Anastasiou for providing the information on protein oxidation assay and Mr. Y.B. Ong for his technical support in the animal study. The work is supported by research grants from Singapore National Medical Research Council (NMRC/CIRG/1373/2013 and NMRC/CIRG/1430/2015) to H.-M.S.. Y.R., J.C., X.M., Q.Y., P.C., G.L. and H.W.-S.T. are supported by research scholarships from the National University of Singapore (NUS).

## AUTHOR CONTRIBUTIONS

Conceptualization, Y.R. and H.-M.S.; Methodology, Y.R., J.C., G.L., H.W.-S.T., Y.J. and Q.Yu; Investigation, Y.R., J.C., X.M., Q.Yang and P.C.; Formal Analysis, Y.R.; Writing – Original Draft, Y.R.; Writing - Review & Editing, H.-M.S., Q.Yu and Y.S.K.; Funding Acquisition, H.-M.S.; Supervision, H.-M.S.

## STAR METHODS

### KEY RESOURCES TABLE

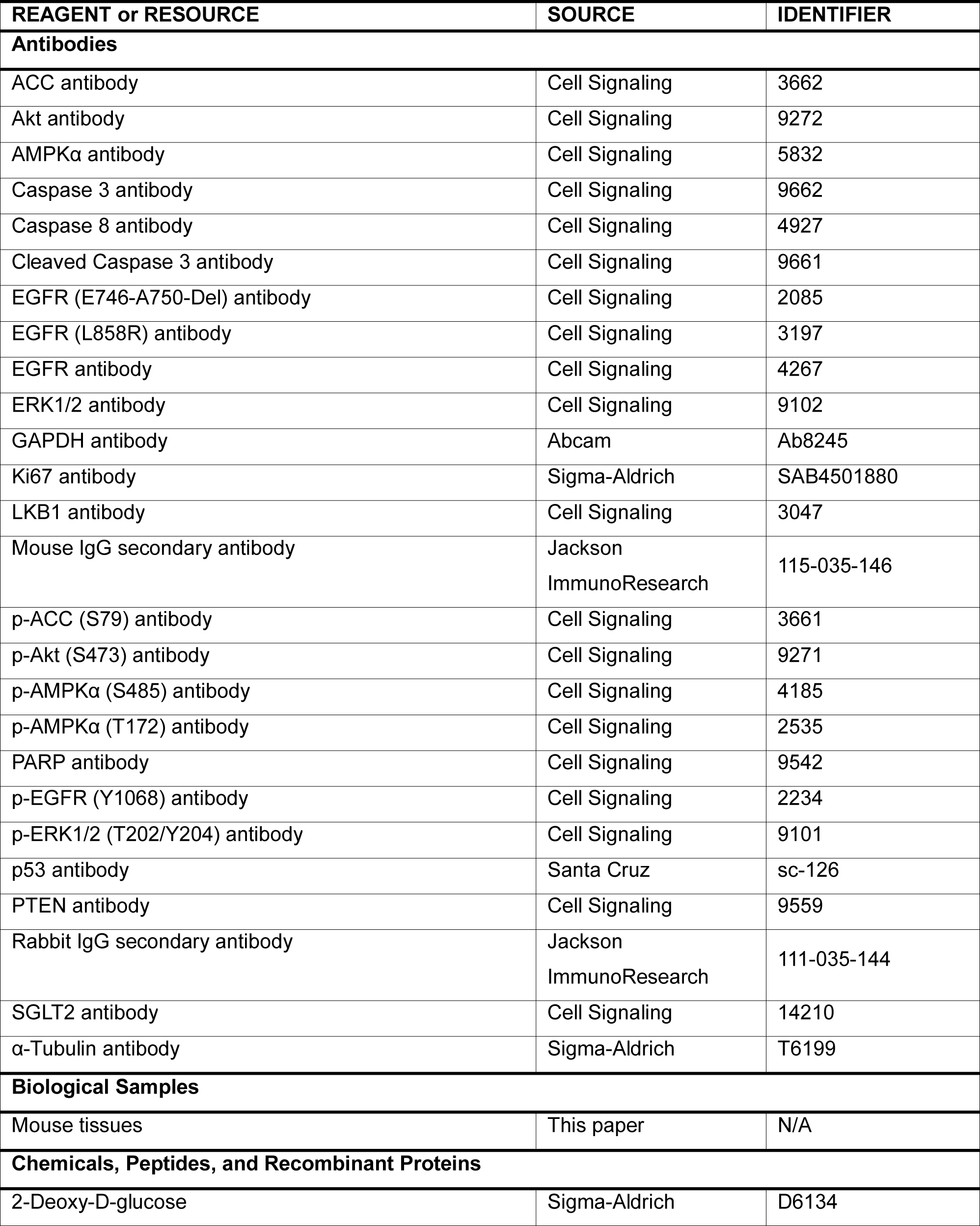
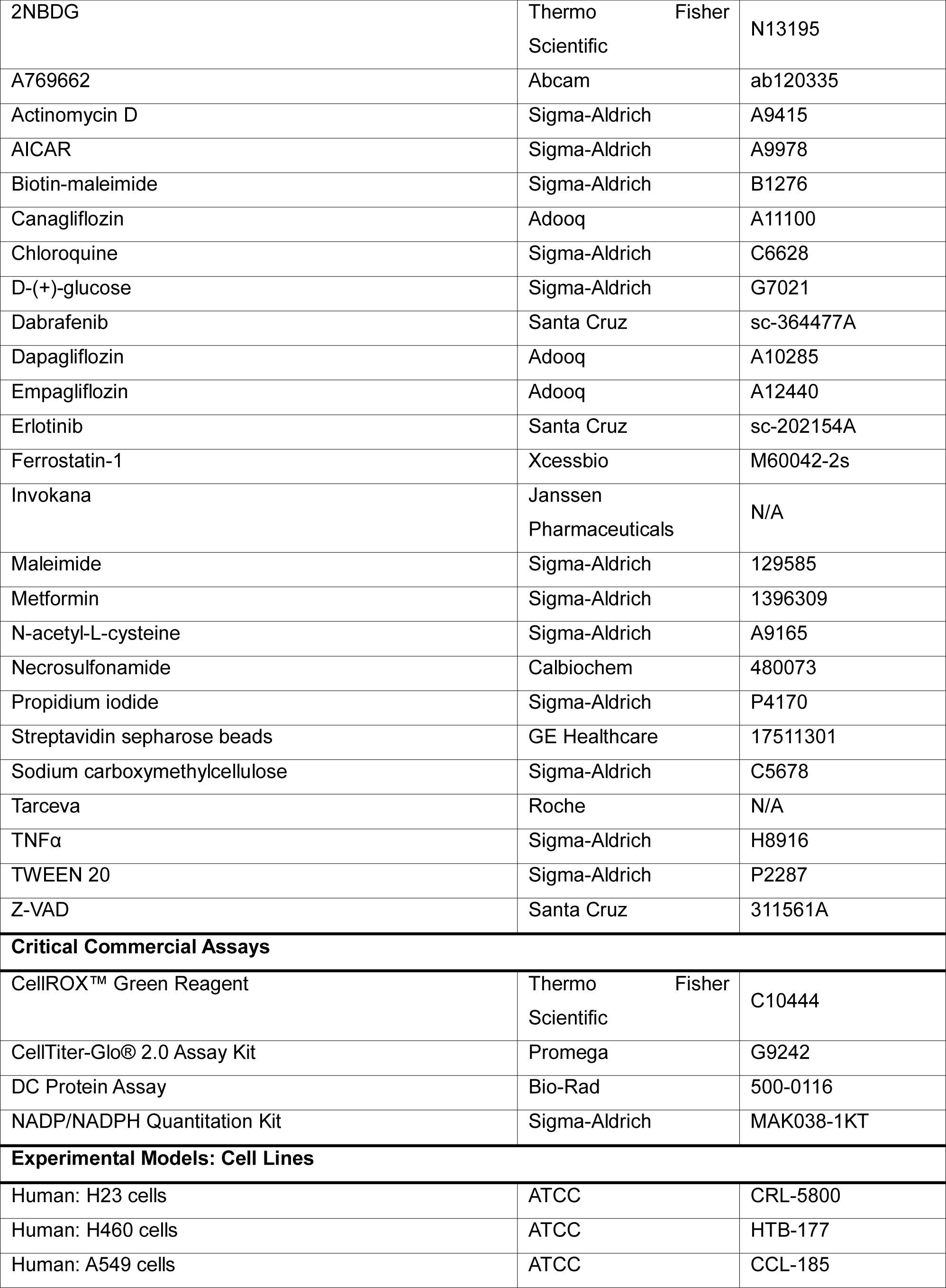
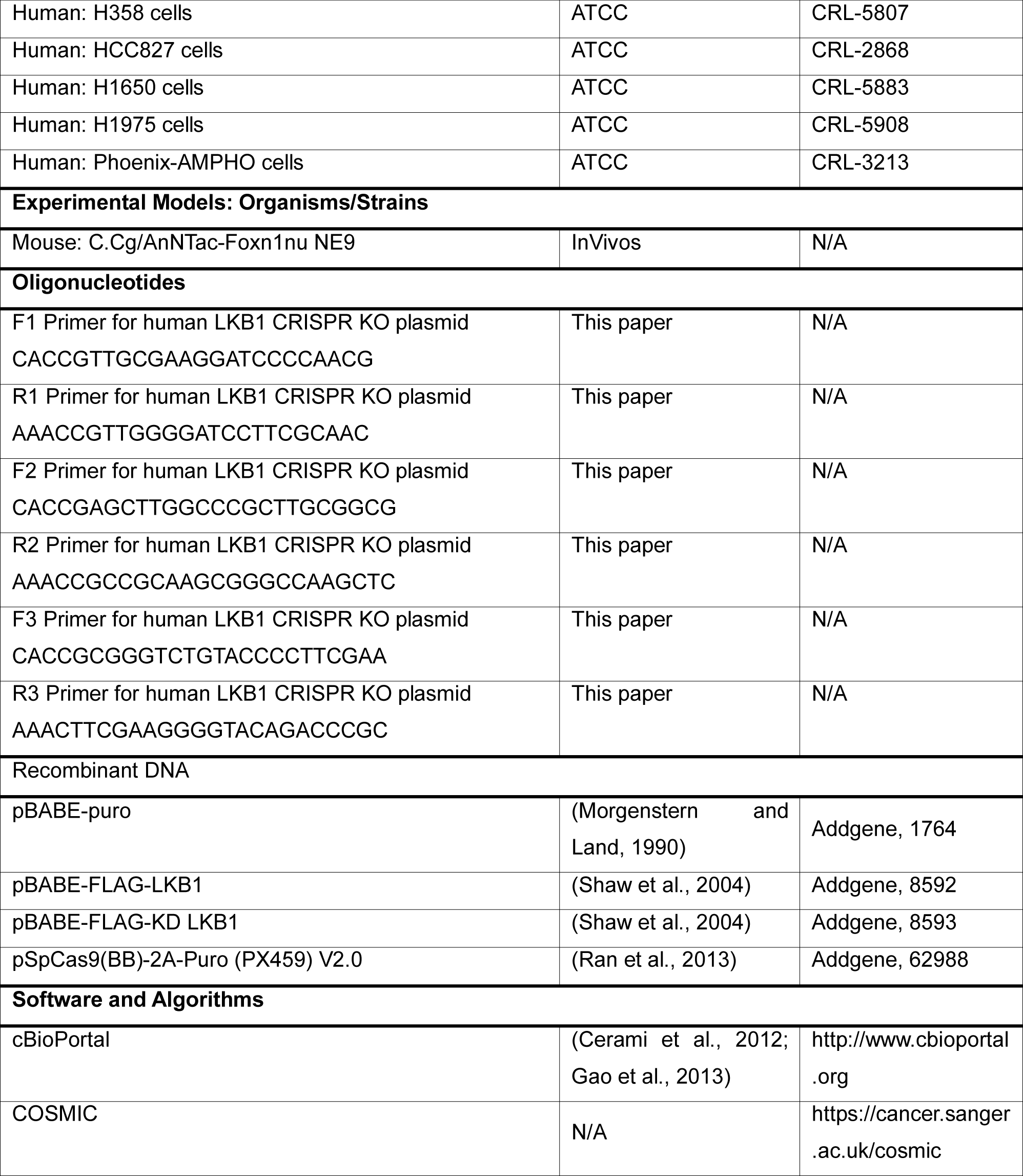

### CONTACT FOR REAGENT AND RESOURCE SHARING

Further information and requests for resources and reagents should be directed to and will be fulfilled by the Lead Contact, Han-Ming Shen

## EXPERIMENTAL MODELS AND SUBJECT DETAILS

### Cell Lines

H460 and A549 cell lines were cultured in Dulbecco’s Modified Eagle Medium (DMEM, Hyclone, SH3002201) supplemented with 10% FBS (Hyclone, SV30160.03). H23, H358, HCC827, H1650 and H1975 cell lines were cultured in Roswell Park Memorial Institute (RPMI) 1640 medium (Thermo Fisher Scientific, A1049101) supplemented with 10% FBS. All cells were maintained in 5% CO_2_ incubator at 37°C.

### Mice

Female 6 to 8 weeks old BALB/c nude mice (C.Cg/AnNTac-Foxn1nu NE9), obtained from InVivos) were used. Animal health was monitored daily by observation and measurement of mice body weight. Animal experiments were conducted according to an approved protocol by the National University of Singapore (NUS) Institutional Animal Care and Use Committee (IACUC) which conforms to NIH guidelines (IACUC protocol # R16-0840). H460 (1 × 10^6^), A549 (1 × 10^6^) and HCC827 (5 × 10^6^) cells were injected subcutaneously into the right flank of the mice. When tumors reached a volume of approximately 50-100 mm^3^, mice were randomized into the following 3 groups: vehicle (0.5% carboxymethyl cellulose and 0.025% Tween-20), Invokana (100 mg/kg/day) or Tarceva (50 mg/kg/day), administered by oral gavage daily. Tumor volume was measured daily after treatments start, based on caliper measurements and calculated by the modified ellipsoidal formula: tumor volume = ½ length × width^2^. After the mice are sacrificed, the tumor samples were excised from mice for the following experiments (1) preparation of tumor homogenate for immunoblotting; and (2) preparation of OCT compound-blocks and slices for immunohistochemistry.

## METHOD DETAILS

### Reagents and Antibodies

All the reagents and antibodies used in this study were obtained as listed in the **Key Resources Table**. All the primary antibodies were 1:1,000 diluted in Tris-buffered saline with 0.1% TWEEN 20 (TBST), supplemented with 5% BAS and 0.1% sodium azide, except that antibodies against GAPDH, α-Tubulin, β-Actin were 1:5,000 diluted, and Ki67 antibody was 1:50 diluted for immunohistochemistry. Mouse and rabbit IgG secondary antibodies were 1:5,000 diluted in TBST supplemented with 5% non-fat milk (Bio-Rad).

### Nutrients Starvation

Glucose-free DMEM (Thermo Fisher Scientific, 11966025), glutamine-free DMEM (Thermo Fisher Scientific, 11960044), and amino acid free-DMEM (recipe provided by Dr. Noboru Mizushima) were supplemented with 10% dialyzed FBS (Thermo Fisher Scientific, 26400044) for different types of starvation. The recipe of amino acid free-DMEM is: NaCl, 6.06 g; NaHCO_3_, 3.7 g; D-glucose, 1 g; KCl, 0.4 g; CaCl_2_.2H_2_O, 0.264 g; MgSO_4_.7H_2_O, 0.2 g; NaH_2_PO_4_.2H_2_O 0.11 g; phenol red 0.015 g; MEM vitamin solution (100X), 40 ml; HEPES (1 M, pH7.5) 15 ml; Fe(NO_3_)_3_ (10 mg/ml), 10 μl, ddH_2_O, to 1 l.

### Propidium Iodide (PI) Exclusion Assay

Cells were seeded in 24-well plates one day before treatments. After designed treatments, the medium and cells were all harvested with trypsin and centrifuged at 500 g for 2 min at 4 °C. The cell pellet was washed with cold PBS and centrifuged again. Then the cells were resuspended in cold PBS containing 5 μg/ml PI and subjected to flow cytometry. Ten thousand cells were analyzed for PI fluorescence intensity (FL3 channel) with FACS Calibur Flow Cytometry (BD Bioscience) using CellQuest software.

### CellTiter-Glo 2.0 Assay

Three thousand cells per well were seeded in 96-well plates one day before treatments. After designed treatments, cellular ATP level was measured using CellTiter-Glo 2.0 Assay Kit following the manufacturer’s instruction. Briefly, after treatments, the cell culture plate was equilibrated at room temperature for approximately 30 minutes and then an equal volume of CellTiter-Glo 2.0 Reagent was added to each well. Shake the plate on an orbital shaker for 2 minutes and incubate at room temperature for 10 minutes. Transfer the contents to an opaque-walled 96-well plate and record luminescence using microplate luminometer (Thermo Fisher Scientific, Luminoskan).

### Immunoblotting

Cells were seeded in 12-well plates one day before treatments. After designed treatments, cell lysates are harvested using SDS lysis buffer (20% glycerol, 2% SDS, 62.5 mM Tris (pH6.8), 2 mM DTT, and 1X phosphatases and proteases inhibitor cocktail). Protein concentration was measured using the DC protein assay method (Bio-Rad) and 25 μg protein samples were loaded and separated by SDS-PAGE and then transferred to PVDF membranes. The membranes were blocked with StartingBlock (PBS) Blocking Buffer (Thermo Fisher Scientific, 37538) at room temperature for 30 mins and incubated with primary antibodies overnight at 4°C. Membranes were washed with TBST three times and then incubated with HRP-conjugated secondary antibodies for 1 hr at room temperature. The chemiluminescence signal was visualized with enhanced chemiluminescence buffer (GE Healthcare, RPN2235) using ImageQuant LAS 500 (GE Healthcare). KODAK Molecular Imaging software was used to analyze the band signal.

### Protein Oxidation

Protein oxidation was measured as previously reported (Anastasiou et al., 2011). The cells were lysed for 15 min in biotin-labeling lysis buffer (BLLB: 50 mM Tris-HCl pH 7.0, 5 mM EDTA, 120 mM NaCl, and 0.5% NP-40) containing protease inhibitors and 100 mM maleimide (Sigma) at 4°C. After the lysates were centrifuged, the supernatant was removed to a fresh tube, and the protein concentration between each group was adjusted to the same level using BLLB. SDS was added to a final concentration of 1%, and the cell lysates were incubated at room temperature for 2 h while rotating. To remove any redundant maleimide, 5 volumes of acetone (−20°C) were added, and the cell lysates were incubated for 20 min. After centrifuged at 20,000 g for 10 min at 4°C, the supernatants were discarded, and the precipitated protein was air-dried. A volume of 200 mL of BLLB containing 1% SDS, 10 mM DTT and 0.1 mM biotin-maleimide (Sigma, stock dissolved in dimethylformamide) was added to resuspend the proteins and to allow the reaction between biotin-maleimide and the reduced, previously oxidized, sulfhydryl groups. A total of 5 volumes of methanol (−20°C) was added to precipitate the proteins. The pellets were resuspended in 500 mL BLLB and then incubated with 10 mL of streptavidin-Sepharose beads (GE Healthcare) while rotating at 4°C for 2 h. The beads were then washed 3 times with BLLB and boiled in SDS-PAGE loading buffer to perform SDS-PAGE analysis.

### Measurement of Intracellular ROS Level

Cells were seeded in 24-well plates one day before treatments. Thirty minutes prior to the end of treatments, CellROX Green Reagent was added at a final concentration of 5 μM and cells were incubated for 30 minutes at 37°C. Then the medium was discarded, and the cells were washed with cold PBS for 3 times. Then the cells were resuspended in cold PBS and subjected to flow cytometry. Ten thousand cells were analyzed for CellROX Green Reagent fluorescence intensity (FL1 channel) with FACS Calibur Flow Cytometry (BD Bioscience) using CellQuest software.

### Measurement of Glucose Uptake

Cells were seeded in 24-well plates one day before treatments. Cells were washed with PBS once and changed to glucose-free DMEM to ensure the maximal glucose uptake and to avoid the disturbance of glucose, and then treated with canagliflozin (100 μM) for 2 hrs. Thirty minutes prior to the end of treatments, 2-(N-(7-Nitrobenz-2-oxa-1,3-diazol-4-yl)Amino)-2-Deoxyglucose (2NBDG) was added at a final concentration of 200 μM and cells were incubated for 30 minutes at 37°C. Then the medium was discarded, and the cells were washed with cold PBS for 3 times. Then the cells were resuspended in cold PBS and subjected to flow cytometry. Ten thousand cells were analyzed for CellROX Green Reagent fluorescence intensity (FL1 channel) with FACS Calibur Flow Cytometry (BD Bioscience) using CellQuest software.

### Ki67 Immunohistochemistry

The NSCLC xenograft tumors were excised and frozen in liquid nitrogen before processing. Tumor samples were mounted in OCT compound and cryosectioned at 8 μm thickness. Slides were dried at 37°C for 30 mins and fixed with 10% neutral buffered formalin for 10 mins at room temperature. Sections were washed with PBS for 5 mins twice and then incubated with 3% H_2_O_2_ for 10 min at room temperature to block the endogenous peroxidase activity. Then samples were washed with PBS twice for 5 mins and blocked with 1% BSA in PBS for 1 hr at room temperature, and then incubated with Ki67 antibody (1:50 dilution in PBS) overnight at 4°C. Samples were washed with PBS three times and then incubated with secondary antibody for 1 hr at room temperature. After incubation, samples were washed with PBS three times and incubated with 3, 3-diaminobenzidine (DAB) for 2 mins at room temperature to develop color. Samples were washed with PBS for three times and counterstained with hematoxylin for 10 mins followed dehydration in 95% ethanol twice for 30 sec each and then 100% ethanol twice for 30 sec each. After air dry, samples were covered with coverslips and visualized using a light microscope.

### Retrovirus Production and Transduction

To produce retrovirus, Phoenix-AMPHO packaging cells were transfected with pBABE empty vector (Morgenstern and Land, 1990) or pBABE-FLAG-LKB1 (WT/KD) (Shaw et al., 2004) vectors using Lipofectamine 3000 reagent (Thermo Fisher Scientific). Retroviral supernatants were collected and filtered at 48, 72 and 96 hrs after transfection and used to infect H23, H460 and A549 cell lines for 6 hrs in the presence of 5 μg/mL polybrene. Cells were then selected and maintained with 1 μg/mL puromycin. LKB1 expression was verified by immunoblotting.

### LKB1 CRISPR Knockout

CRISPR sgRNA oligonucleotides targeting human LKB1 gene were designed at http://crispr.mit.edu and then cloned into pSpCas9(BB)-2A-Puro (PX459) V2.0 vector following the protocol (Ran et al., 2013). The vectors were transfected into H358 and HCC827 cell lines using Lipofectamine 3000 reagent (Thermo Fisher Scientific). Monoclones were then selected and maintained with 1 μg/mL puromycin. Positive clones with LKB1 gene knockout was verified by immunoblotting.

### Statistical Analyses

All image data and immunoblotting data presented below were representatives from at least three independent repeated experiments. All numeric data were presented as Mean ± SE from at least 3 independent experiments. The p-value was analyzed by Student’s t test or Two-way ANOVA, and *p < 0.05, **p < 0.01, ***p < 0.001, ****p < 0.001 was considered as statistically significant.

