## Supplemental Figures & Legends for "Synthetic lethality targeting LKB1 mutant and EGFR wild type human non-small cell lung cancer cells by glucose starvation and SGLT2 inhibition"

A

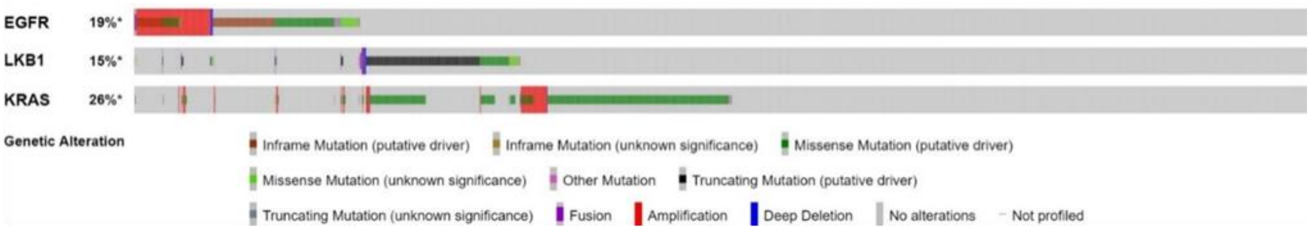


B

| A | B | Neither | A Not B | B Not A | Both | Log Odds Ratio | p-Value | Adjusted p-Value ▲ | Tendency |
|---|---|---|---|---|---|---|---|---|---|
| EGFR | KRAS | 1738 | 575 | 776 | 40 | -1.859 | <0.001 | **<0.001** | Mutual exclusivity Significant |
| LKB1 | KRAS | 2084 | 229 | 589 | 227 | 1.255 | <0.001 | **<0.001** | Co-occurrence Significant |

C

| No | Reference | Cancer Type | Sample Size | Mutation Rate | Mutual Exclusivity Test |
|---|---|---|---|---|---|
| 1 | Jordan E J, et al., Cancer Discov, 2017 | ADC | 915 | EGFR: 30%<br>LKB1: 18% | Significant<br>p-Value: <0.001 |
| 2 | Campbell J D, et al., Nat Genet, 2016 | ADC<br>SqCC | 1144 | EGFR: 14%<br>LKB1: 10% | Significant<br>p-Value: 0.002 |
| 3 | Ding L, et al., Nature, 2008 | ADC | 163 | EGFR: 19%<br>LKB1: 21% | Significant<br>p-Value: 0.004 |
| 4 | Cancer Genome Atlas Research N, Nature, 2014 | ADC | 230 | EGFR: 17%<br>LKB1: 19% | Significant<br>p-Value: 0.031 |
| 5 | Rizvi H, et al., J Clin Oncol, 2018 | NSCLC | 240 | EGFR: 14%<br>LKB1: 23% | Tendency towards mutual exclusivity<br>p-Value: 0.073 |
| 6 | Imielinski M, et al., Cell, 2012 | ADC | 183 | EGFR: 19%<br>LKB1: 15% | Tendency towards mutual exclusivity<br>p-Value: 0.183 |
| 7 | Rizvi N A, et al., Science, 2015 | ADC | 35 | EGFR: 6%<br>LKB1: 15% | Tendency towards mutual exclusivity<br>p-Value: 0.724 |
| 8 | Cancer Genome Atlas Research N, Nature, 2012 | SqCC | 178 | EGFR: 9%<br>LKB1: 1.7% | Tendency towards mutual exclusivity<br>p-Value: 0.753 |
| 9 | Vavala T, et al., Lung Cancer, 2017 | NSCLC | 41 | EGFR: 29%<br>LKB1: 12% | Tendency towards co-occurrence<br>p-Value: 0.461 |

**Figure S1. EGFR and LKB1 Mutations Are Mutually Exclusive in NSCLC. Related to Figure 1.**

**(A)** KRAS mutation is included in the mutation spectrum of NSCLC. The same set of samples in **Figure 1A** were pooled and analyzed using the cBioPortal database.

**(B)** The mutations of EGFR and LKB1/KRAS are mutually exclusive in NSCLC samples. Same sets of samples in (**A**) were tested for their mutual exclusivity using the cBioPortal database.

**(C)** The mutation rate and the mutual exclusivity test of 9 NSCLC studies are listed.

ADC, adenocarcinoma; SqCC, squamous cell carcinoma; NSCLC, non-small cell lung cancer. All data were obtained from cBioPortal (http://www.cbioportal.org).

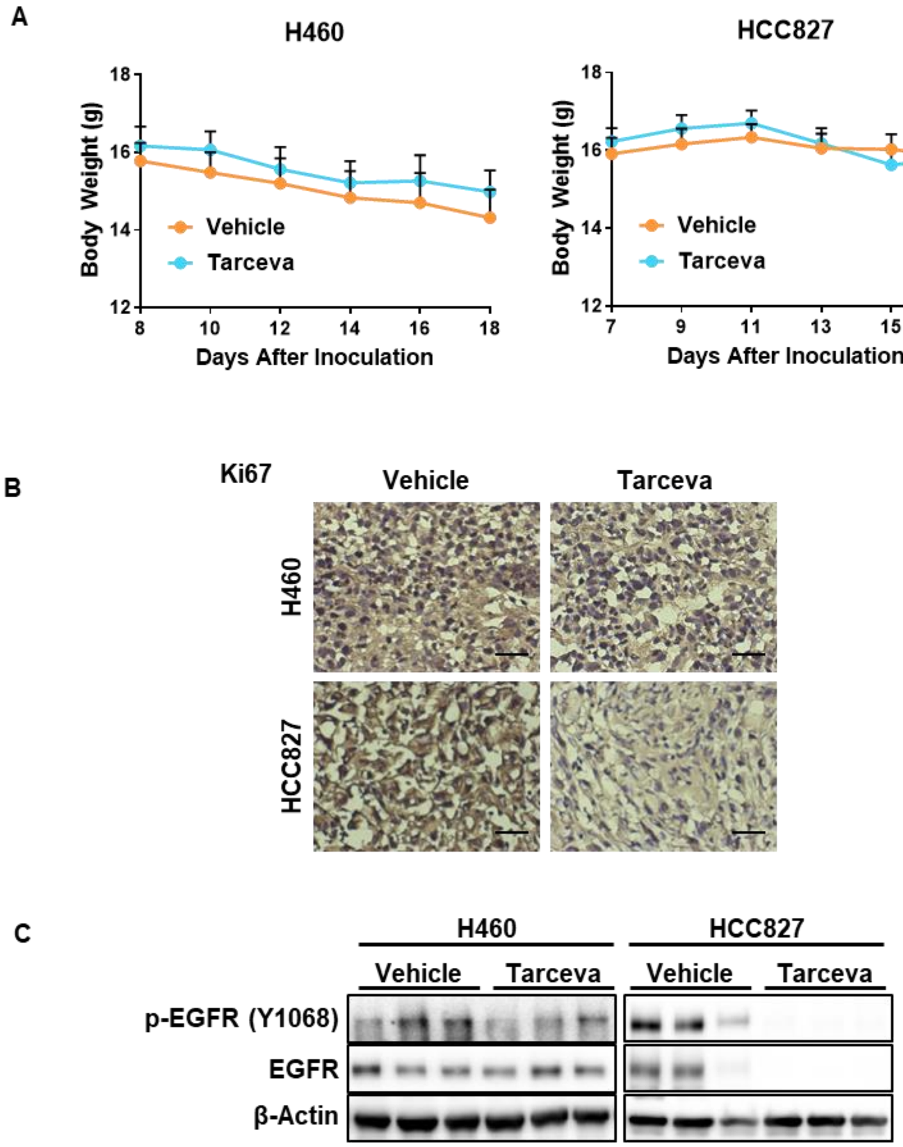
A
H460
HCC827
Body Weight (g)
18
16
14
12
Vehicle
Tarceva
8
10
12
14
16
18
7
9
11
13
15
17
Days After Inoculation
B
Ki67
Vehicle
Tarceva
H460
HCC827
C
H460
HCC827
Vehicle
Tarceva
p-EGFR (Y1068)
EGFR
β-Actin

**Figure S2. LKB1-Mutant/EGFR-WT NSCLC Cells Are Resistant to EGFR Inhibitor Erlotinib. Related to Figure 2.**

**(A)** Tarceva has marginal side effects on mice body weight. Mice body weight was recorded together with tumor volume. Data were presented as Mean ± SE from 6 (H460) or 8 (HCC827) mice per group. (two-way ANOVA).

**(B)** Tarceva exhibits weaker effects on inhibiting cell proliferation of LKB1-mutant NSCLC cells. Tumors formed by H460 and HCC827 cell lines were excised from mice 24 hrs after the last gavage. Samples were embedded, frozen, sectioned and stained for cell proliferation marker Ki67. Representative images are shown. Scale bar, 100 μm.

**(C)** Tarceva has marginal effects on EGFR signaling pathway in tumors formed by LKB1-mutant NSCLC cells. Tumors formed by H460 and HCC827 cell lines were excised from mice 24 hrs after the last gavage. Samples were lysed for immunoblotting.

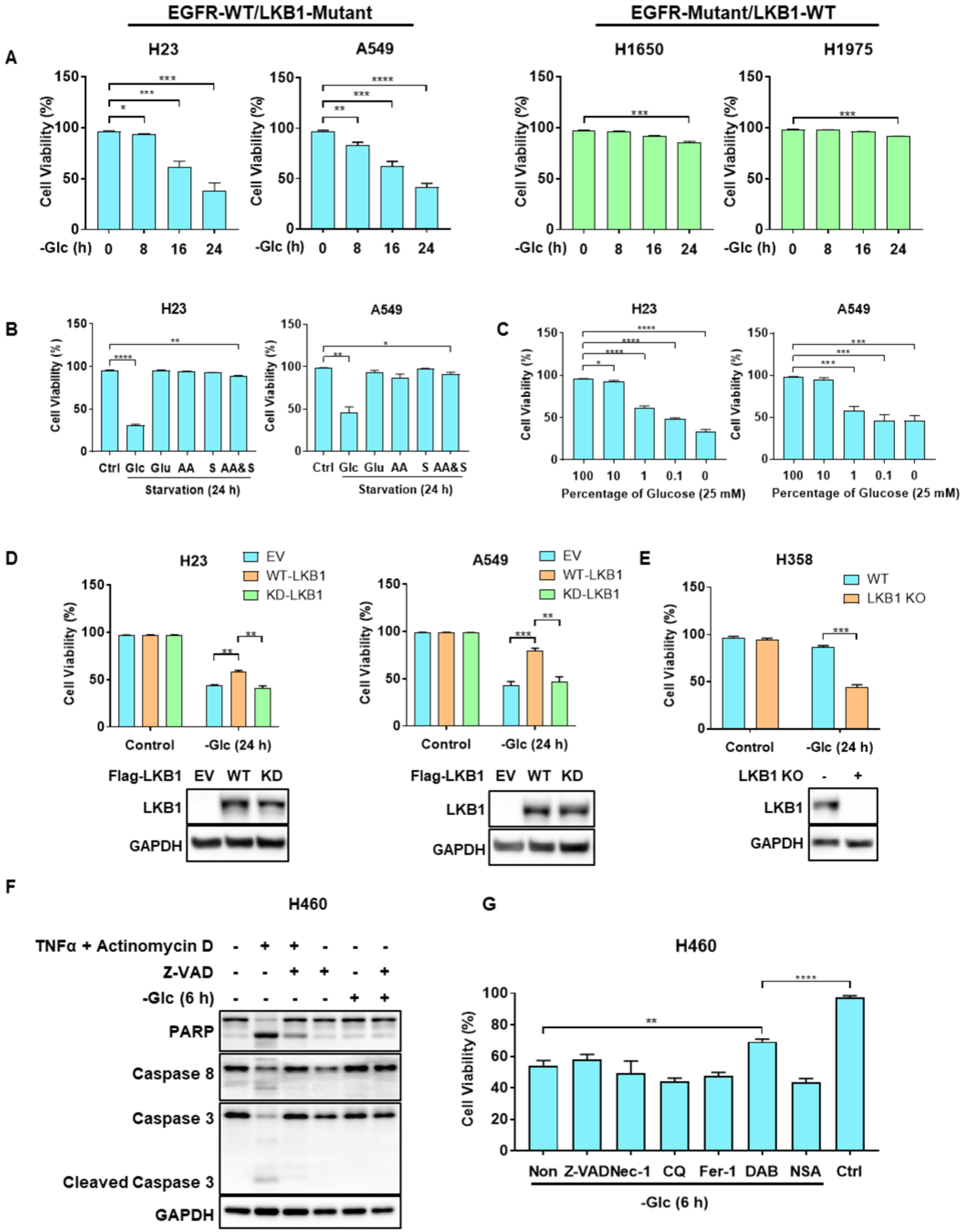

EGFR-WT/LKB1-Mutant
EGFR-Mutant/LKB1-WT
A
H23
A549
H1650
H1975
Cell Viability (%)
-Glc (h) 0 8 16 24
B
Ctrl Glc Glu AA S AA&S
Starvation (24 h)
C
Percentage of Glucose (25 mM)
100 10 1 0.1 0
D
EV
WT-LKB1
KD-LKB1
Control
-Glc (24 h)
Flag-LKB1 EV WT KD
LKB1
GAPDH
E
H358
WT
LKB1 KO
LKB1 KO - +
F
H460
TNFα + Actinomycin D
Z-VAD
-Glc (6 h)
PARP
Caspase 8
Caspase 3
Cleaved Caspase 3
GAPDH
G
H460
Non Z-VAD Nec-1 CQ Fer-1 DAB NSA Ctrl
-Glc (6 h)

**Figure S3. LKB1-Mutant/EGFR-WT NSCLC Cells Are Sensitive to Cell Death Induced by Glucose Starvation. Related to Figure 3.**

**(A)** LKB1-mutant NSCLC cell lines are more sensitive to glucose starvation. The other 4 NSCLC cell lines were treated with glucose starvation for the indicated times. Cell viability was measured by PI exclusion assay using flow cytometry.

**(B)** LKB1-mutant NSCLC cell lines are only sensitive to glucose starvation. H23 and A549 cell lines were cultured in DMEM without glucose, glutamine, amino acids (AA), serum, or the combination of AA and serum for 24 hrs. Cell viability was measured by PI exclusion assay using flow cytometry.

**(C)** Glucose replenishment rescues cell death in a dose-dependent manner in LKB1-mutant NSCLC cell lines. H23 and A549 cell lines were cultured in glucose-free DMEM, supplemented with the indicated percentage of glucose for 24 hrs. Cell viability was measured by PI exclusion assay using flow cytometry.

**(D)** LKB1 reconstruction activates AMPK and rescues cell death in LKB1-mutant NSCLC cell lines upon glucose starvation. H23 and A549 cell lines stably expressing empty vector (EV), wild type (WT)- or kinase dead (KD)-LKB1 were treated with full or glucose-free DMEM for 24 hrs. Cell viability was measured by PI exclusion assay using flow cytometry. LKB1 reconstitution was shown.

**(E)** LKB1 knockout sensitizes LKB1-WT NSCLC cell lines to cell death induced by glucose starvation. Wild type (WT) or LKB1 knockout (KO) H358 cell line was treated with full or glucose-free DMEM for 24 hrs. Cell viability was measured by PI exclusion assay using flow cytometry. LKB1 knockout was shown.

**(F)** Glucose starvation induces non-apoptotic cell death in LKB1-mutant H460 cells. H460 cell line cultured in full or glucose-free DMEM were treated with Z-VAD (40 μM) for 6 hrs, TNFα and Actinomycin D were used as an apoptosis inducer for the positive control. Cells were lyzed and cleavage of PARP, caspase 3 and caspase 8 were analyzed by immunoblotting.

**(G)** Glucose starvation induces non-apoptotic, non-necroptotic, non-autophagic and non-ferroptotic cell death in LKB1-mutant H460 cells. H460 cell line cultured in glucose-free DMEM were treated with Z-VAD (40 μM), necrostatin-1 (Nec-1) (30 μM), chloroquine (CQ) (50 μM), ferrostatin-1 (Fer-1) (5 μM), dabrafenib (DAB) (20 μM), or necrosulfonamide (NSA) (2 μM) for 6 hrs. Cell viability was measured by PI exclusion assay using flow cytometry.

Data were presented as Mean ± SE from at least 3 independent experiments. *p < 0.05, **p < 0.01, ***p < 0.001, ****p < 0.0001 (student's t test) comparing to their respective control (full medium or non-treated) group.

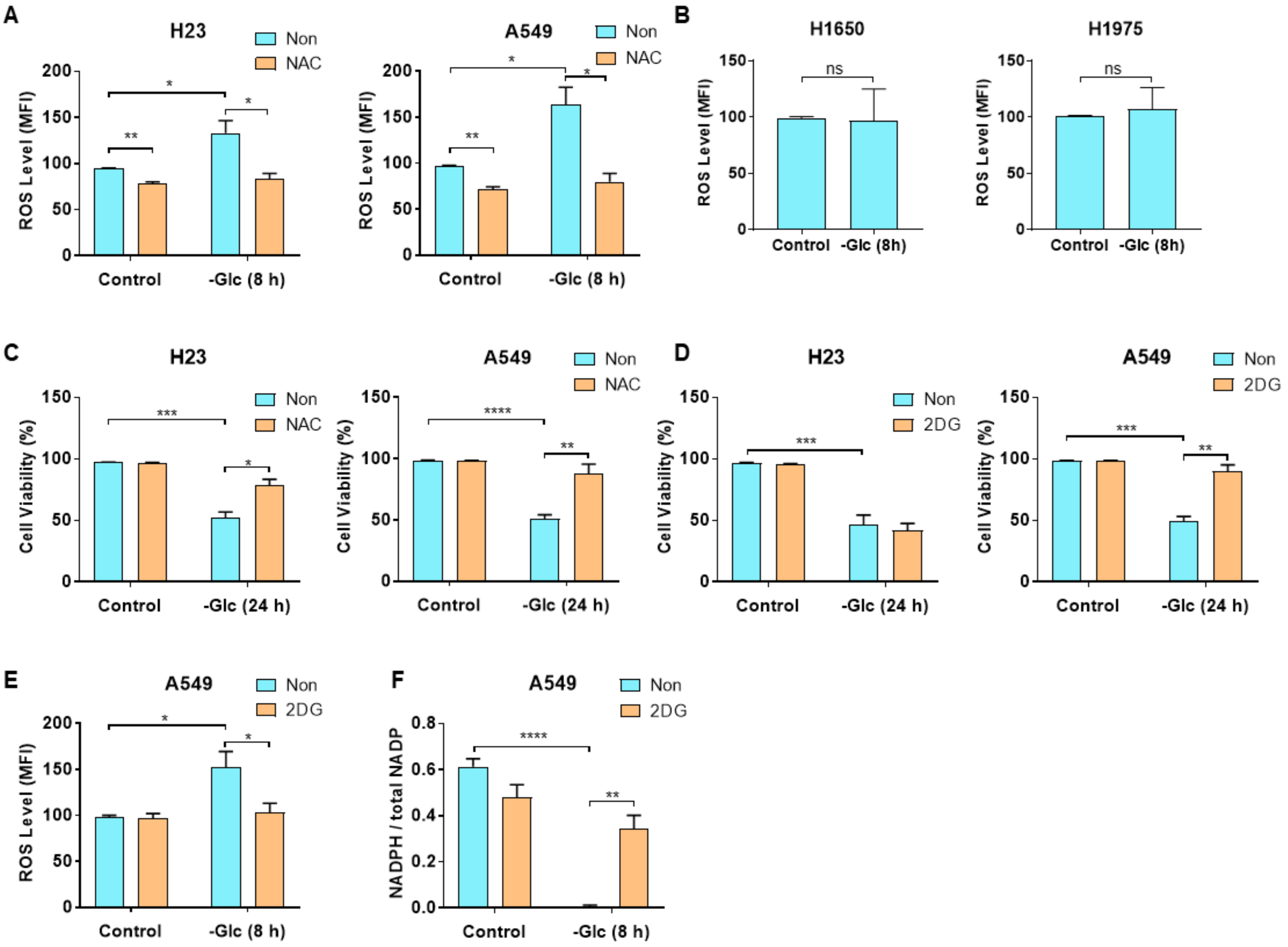
A
H23
Non
NAC
ROS Level (MFI)
200
150
100
50
0
Control
-Glc (8 h)
**
*
*
A549
B
H1650
H1975
ns
-Glc (8h)
C
Cell Viability (%)
-Glc (24 h)
***
****
D
2DG
E
F
NADPH / total NADP
0.8
0.6
0.4
0.2
0.0

**Figure S4. Glucose Starvation Induces Cell Death via Oxidative Stress in EGFR-WT/LKB1-Mutant NSCLC Cells. Related to Figure 4.**

**(A and B)** Glucose starvation increases the ROS level in LKB1-mutant NSCLC cells. **(A)** H23 and A549 cell lines cultured in full or glucose-free DMEM were treated with NAC (5 mM) for 8 hrs. **(B)** H1650 and H1975 cell lines were treated with full or glucose-free DMEM for 8 hrs. Intracellular ROS levels were measured using CellROX Green reagent by flow cytometry.

**(C)** NAC blocks glucose starvation-induced cell death in LKB1-mutant NSCLC cell lines H23 and A549 cell lines cultured in full or glucose-free DMEM were treated with NAC (5 mM) for 8 hrs. Cell viability was measured by PI exclusion assay using flow cytometry.

**(D)** 2DG rescues glucose starvation-induced cell death in some LKB1-mutant NSCLC cell lines. H23 and A549 cell lines cultured in full or glucose-free DMEM were treated with 2DG (5 mM) for 8 hrs. Cell viability was measured by PI exclusion assay using flow cytometry.

**(E)** 2DG reduces ROS level in LKB1-mutant NSCLC cell lines under glucose starvation. A549 cell line cultured in full or glucose-free DMEM were treated with 2DG (5 mM) for 8 hrs. Intracellular ROS levels were measured using CellROX Green reagent by flow cytometry.

**(F)** 2DG maintains NAPDH level in LKB1-mutant NSCLC cell lines under glucose starvation. A549 cell line cultured in full or glucose-free DMEM were treated with 2DG (5 mM) for 8 hrs. NADPH and NADP levels were measured using NADP/NADPH Quantitation Kit following the manufacturer's instruction.

Data were presented as Mean ± SE from at least 3 independent experiments. ns, not significant, *p < 0.05, **p < 0.01, ***p < 0.001, ****p < 0.0001 (student's t test) comparing to their respective control (full medium or non-treated) group.

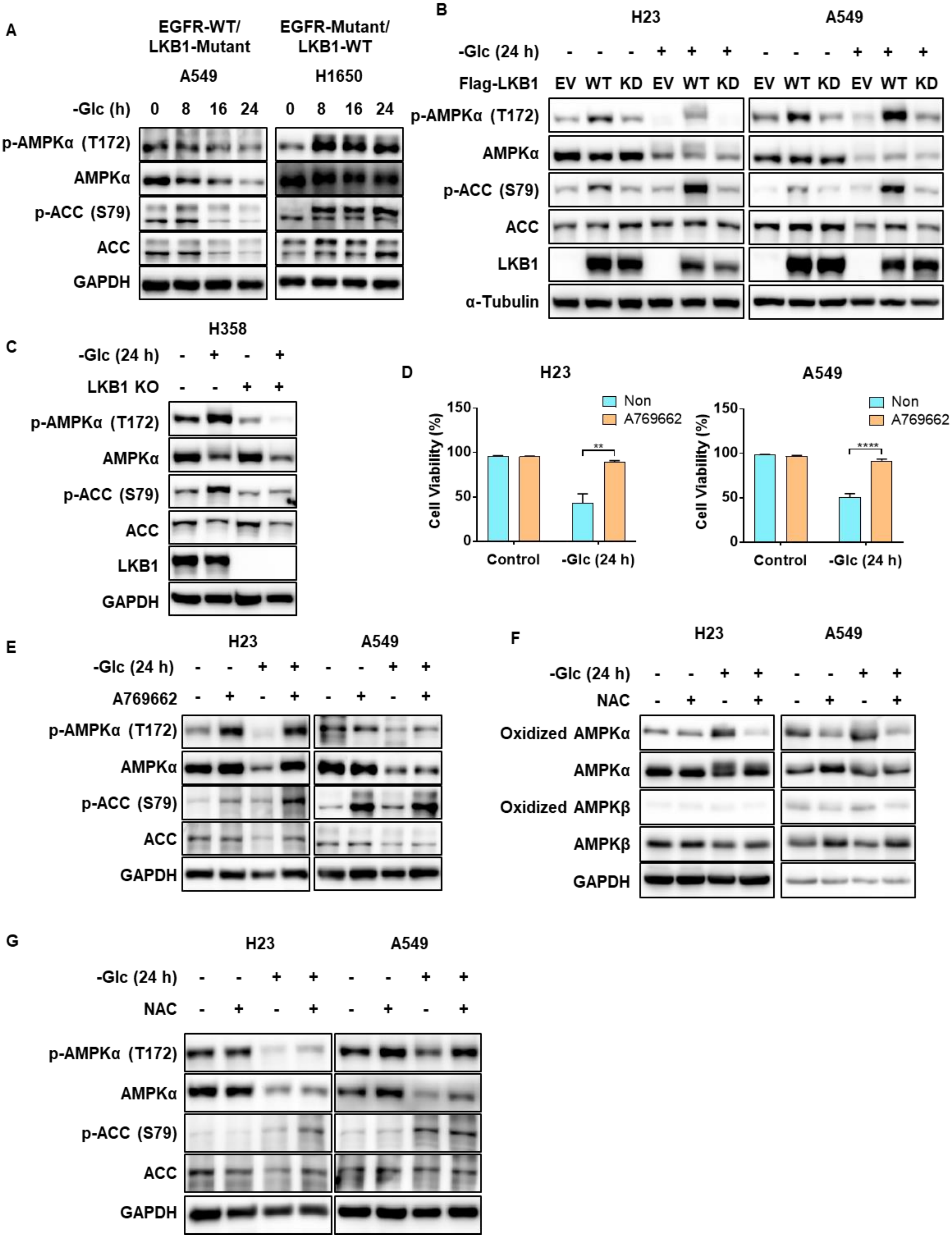
A
EGFR-WT/ LKB1-Mutant
EGFR-Mutant/ LKB1-WT
A549
H1650
-Glc (h) 0 8 16 24 0 8 16 24
p-AMPKα (T172)
AMPKα
p-ACC (S79)
ACC
GAPDH
B
H23
A549
-Glc (24 h) - - - + + + - - - + + +
Flag-LKB1 EV WT KD EV WT KD EV WT KD EV WT KD
p-AMPKα (T172)
AMPKα
p-ACC (S79)
ACC
LKB1
α-Tubulin
C
H358
-Glc (24 h) - + - +
LKB1 KO - - + +
p-AMPKα (T172)
AMPKα
p-ACC (S79)
ACC
LKB1
GAPDH
D
H23
A549
Non
A769662
Cell Viability (%)
150
100
50
0
**
****
Control
-Glc (24 h)
E
H23
A549
-Glc (24 h) - - + + - - + +
A769662 - + - + - + - +
p-AMPKα (T172)
AMPKα
p-ACC (S79)
ACC
GAPDH
F
H23
A549
-Glc (24 h) - - + + - - + +
NAC - + - + - + - +
Oxidized AMPKα
AMPKα
Oxidized AMPKβ
AMPKβ
GAPDH
G
H23
A549
-Glc (24 h) - - + + - - + +
NAC - + - + - + - +
p-AMPKα (T172)
AMPKα
p-ACC (S79)
ACC
GAPDH

**Figure S5. Impaired AMPK Activation in EGFR-WT/LKB1-Mutant NSCLC Cells Is Implicated in The Susceptibility to Cell Death under Glucose Starvation. Related to Figure 5.**

**(A)** AMPK is inactivated in LKB1-mutant NSCLC cell lines upon glucose starvation. H23 and H1650 cell lines were treated with glucose-free DMEM for the indicated times and then lysed. AMPK activity was measured by ACC and AMPKα phosphorylation using immunoblotting.

**(B)** LKB1 reconstruction activates AMPK in LKB1-mutant NSCLC cell lines upon glucose starvation. H23 and A549 cell lines stably expressing empty vector (EV), wild type (WT)- or kinase dead (KD)-LKB1 were treated with full or glucose-free DMEM for 24 hrs. AMPK activity was measured by ACC and AMPKα phosphorylation using immunoblotting.

**(C)** LKB1 knockout impairs AMPK activation and sensitizes LKB1-WT NSCLC cell lines to cell death induced by glucose starvation. Wild type (WT) or LKB1 knockout (KO) H358 cell line was treated with full or glucose-free DMEM for 24 hrs. AMPK activity was measured by ACC and AMPKα phosphorylation using immunoblotting.

**(D)** Only AMPK allosteric activator A769662 can block cell death in LKB1-mutant NSCLC cell lines upon glucose starvation. H23 and A549 cell lines cultured in full or glucose-free DMEM were treated with A769662 (200 μM) for 24 hrs. Cell viability was measured by PI exclusion assay using flow cytometry. Data were presented as Mean ± SE from at least 3 independent experiments. $^{**}p < 0.01$, $^{****}p < 0.0001$ (student's t test) comparing to non-treated group.

**(E)** AMPK allosteric activator A769662 activates AMPK in LKB1-mutant NSCLC cell lines upon glucose starvation. H23 and A549 cell lines as treated in **(D)** were lysed and AMPK activity was measured by ACC and AMPKα phosphorylation using immunoblotting.

**(F)** AMPKα is oxidized in LKB1-mutant NSCLC cell lines upon glucose starvation. H23 and A549 cell lines cultured in full or glucose-free DMEM were treated with NAC (5 mM) for 24 hrs. Cysteine oxidation of AMPK subunits was detected.

**(G)** NAC maintains AMPK activity in LKB1-mutant NSCLC cell lines upon glucose starvation. H23 and A549 cell lines as treated in **(F)** were lysed and AMPK activity was measured by ACC and AMPKα phosphorylation using immunoblotting.

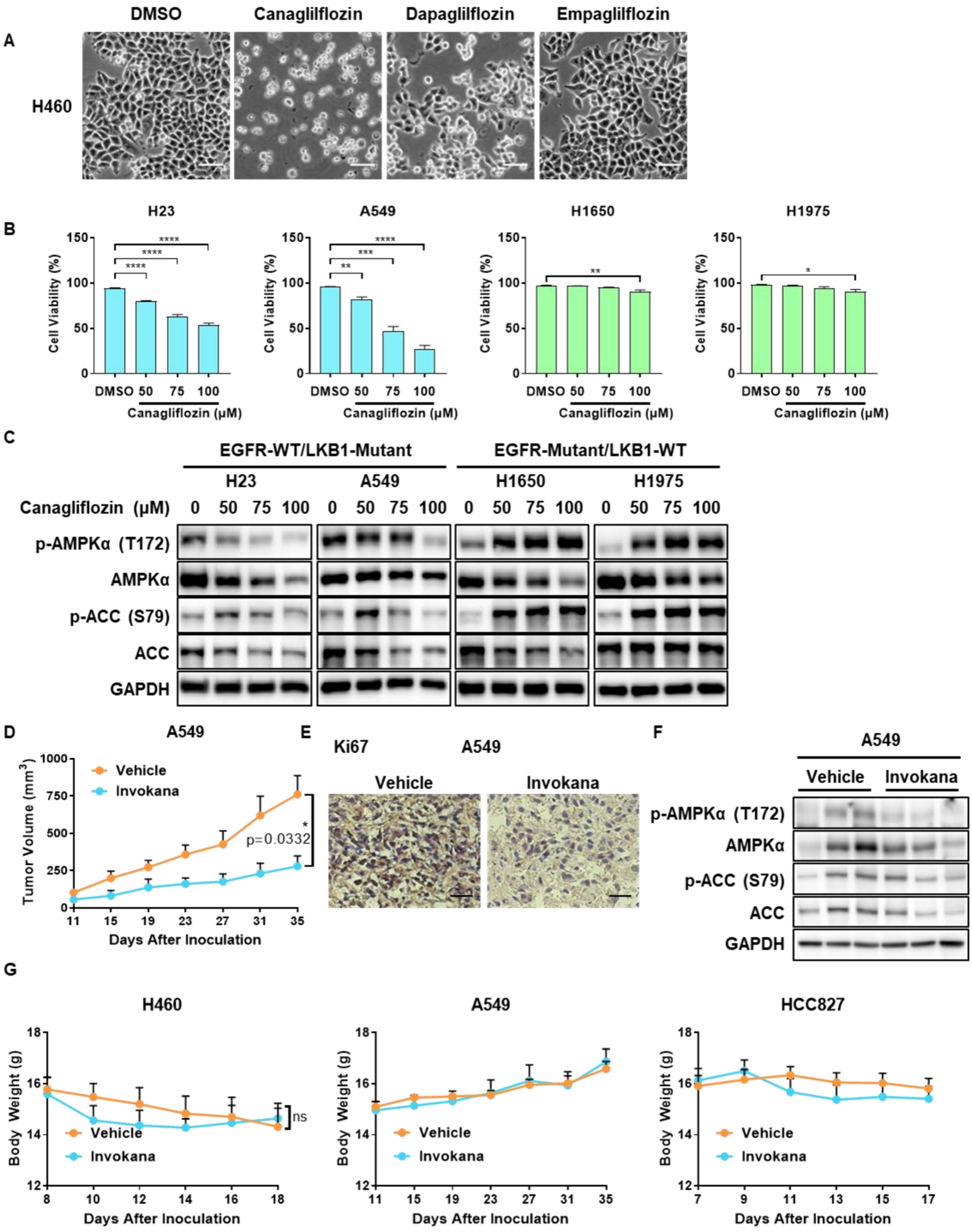

A
DMSO
Canaglilflozin
Dapaglilflozin
Empaglilflozin
H460
B
H23
A549
H1650
H1975
Cell Viability (%)
DMSO 50 75 100
Canagliflozin (μM)
C
EGFR-WT/LKB1-Mutant
EGFR-Mutant/LKB1-WT
H23
A549
H1650
H1975
Canagliflozin (μM) 0 50 75 100 0 50 75 100 0 50 75 100 0 50 75 100
p-AMPKα (T172)
AMPKα
p-ACC (S79)
ACC
GAPDH
D
A549
Vehicle
Invokana
Tumor Volume (mm3)
p=0.0332
Days After Inoculation
E
Ki67
A549
Vehicle
Invokana
F
A549
Vehicle
Invokana
p-AMPKα (T172)
AMPKα
p-ACC (S79)
ACC
GAPDH
G
H460
A549
HCC827
Body Weight (g)
ns
Vehicle
Invokana
Days After Inoculation

**Figure S6. SGLT2 Inhibitor Canagliflozin Causes Synthetic Lethality in LKB1-Mutant NSCLC Cells. Related to Figure 6.**

**(A)** Only canagliflozin can cause cell death in LKB1-mutant NSCLC cell lines. H460 cell line cultured in low glucose DMEM (1 g/L) was treated with 100 μM canagliflozin, dapagliflozin or empagliflozin for 24 hrs. Cell viability was monitored by cell morphology. Representative images were shown. Scale bar, 100 μm.

**(B)** LKB1-mutant NSCLC cell lines are sensitive to canagliflozin. Other 4 NSCLC cell lines cultured in low glucose DMEM (1 g/L) were treated with indicated concentrations of canagliflozin for 24 hrs. Cell viability was measured by PI exclusion assay using flow cytometry.

**(C)** AMPK is inactivated in LKB1-mutant NSCLC cell lines under canagliflozin treatment. Other 4 NSCLC cell lines as treated in **(B)** were lysed. AMPK activity was measured by ACC and AMPKα phosphorylation using immunoblotting.

**(D and E)** Invokana significantly mitigates tumor growth and cell proliferation of LKB1-mutant NSCLC cells *in vivo*. A549 cell line was injected into the flank of BALB/c nude mice. After 11 days when the palpable tumor was established (50 $mm^3$), mice were administered with vehicle or Invokana (100 mg/kg/day) by oral gavage daily. **(D)** Tumor volume was recorded daily. Tumors were excised from mice 24 hrs after the last gavage. Samples were embedded, frozen, sectioned and stained for cell proliferation marker Ki67. **(E)** Representative images are shown. Data were presented as Mean ± SE from at least 6 mice per group. ns, not significant; ***$p < 0.001$ (two-way ANOVA).

**(F)** Invokana only activates AMPK in tumors formed by LKB1-WT NSCLC cells. Tumors formed by A549 cell line were excised from mice 24 hrs after the last gavage. Samples were lysed and AMPK activity was measured by ACC and AMPKα phosphorylation using immunoblotting.

**(G)** Invokana has marginal side effects on mice body weight. Mice body weight was recorded together with tumor volume. Data were presented as Mean ± SE from at least 6 mice per group. ns, not significant (two-way ANOVA).
